# Liquid-liquid phase separation facilitates the biogenesis of secretory storage granules

**DOI:** 10.1101/2021.12.22.472607

**Authors:** Anup Parchure, Meng Tian, Cierra K Boyer, Shelby C Bearrows, Kristen E Rohli, Jianchao Zhang, Bulat R Ramazanov, Yanzhuang Wang, Samuel B Stephens, Julia von Blume

**Affiliations:** Department of Cell Biology, Yale University School of Medicine, New Haven, CT, USA; Fraternal Order of Eagles Diabetes Research Center, Departments of Pharmacology and Neuroscience and Internal Medicine; Division of Endocrinology and Metabolism, University of Iowa, Iowa City, IA, USA; Department of Molecular, Cellular and Developmental Biology, University of Michigan, Ann Arbor, MI, USA; Department of Neurology, University of Michigan School of Medicine, Ann Arbor, MI, USA

**Keywords:** Proinsulin, sorting and secretion, regulated secretion, trans-Golgi Network, LLPS, pH, zinc, calcium, secretory granule biosynthesis

## Abstract

Insulin is a key regulator of human metabolism, and its dysfunction leads to diseases such as type 2 diabetes. It remains unknown how proinsulin is targeted from the trans-Golgi network (TGN) to secretory storage granules as no cargo receptor has been identified. Chromogranin proteins (CGs) are central regulators of granule biosynthesis, and it was proposed that their aggregation is critical for this process. However, the molecular mechanism by which these molecules facilitate sorting at the TGN is poorly understood. Here, we show that CGs undergo liquid–liquid phase separation (LLPS) at low pH independently of divalent cations, such as calcium. Liquid CG condensates, but not aggregates, recruit and sort proinsulin and other granule-destined cargo molecules towards secretory granules. Cargo selectivity is independent of sequence or structural elements but is based on the size and concentration of the client molecules at the TGN. Finally, electrostatic interactions and the N-terminal intrinsically disordered domain of chromogranin B facilitate LLPS and are critical for granule formation. We propose that phase-separated CGs act as a “cargo sponge” within the TGN lumen, gathering soluble client proteins into the condensate independently of specific sequence or structural elements, facilitating receptor-independent sorting. These findings challenge the canonical TGN sorting models and provide insights into granule biosynthesis in insulin-secreting β-cells.

**One sentence summary:** Liquid Chromogranin condensates recruit cargo molecules at the TGN for their delivery to secretory storage granules.

## Introduction

Secretory proteins control human metabolism and physiology, and their mistargeting causes several diseases, including type 2 diabetes (T2D), neurological disorders, and cancer (*1, 2*). Secretory proteins are synthesized in the endoplasmic reticulum (ER) before being loaded into vesicular carriers for transport to the Golgi apparatus (*3–6*). Eventually, these proteins reach the trans-Golgi network (TGN) and are sorted and packaged into specific carriers for transport to their final destinations, including the cell surface, endolysosomes, and secretory storage granules (*7*– *10*). Cargo receptors targeting soluble proteins to the cell surface or secretory granules in specialized cells have not been identified and it remains an enigma how these non-membrane attached molecules are sorted and targeted in the TGN (*9, 11*).

Understanding the sorting and targeting of proteins from the TGN to secretory storage granules in professional secretory cells such as insulin-secreting pancreatic β islets remains a challenge in the field (*12–15*). Secretory storage granules are cytosolic storage structures found in endocrine, exocrine, and neuronal cells. Early electron microscopy imaging has shown that secretory granules contain an electron-dense core, and that condensation occurs in sequential steps starting in the TGN (*16–18*). Soluble proinsulin is synthesized in the ER and transported to the TGN to be sorted and packaged into immature secretory storage granules (ISGs) (*19, 20*). In ISGs, proinsulin is proteolyzed to mature insulin by the protein convertases CPE, PC1/3 and PC2 (*21*), after which mature insulin forms hexameric crystals in the presence of zinc (*21, 22*). In response to nutrient stimuli, mature SGs are mobilized to fuse with the plasma membrane and deliver insulin to the bloodstream, a process called regulated secretion (*23*). While there is considerable knowledge of the processing of proinsulin in SGs, it is unclear how proinsulin is sorted into secretory granules at the TGN (*24*). Importantly, morphometric studies of human T2D islets have shown that the volume density of total mature insulin granules in β-cells is significantly diminished, and that this pool size correlates with the diminished magnitude of the glucose-stimulated insulin secretion response (*25*).

Pulse-chase experiments have shown that dense core structures emerge from the Golgi to form ISGs and that granule content further condenses upon reaching the mature secretory granule (*18, 26*). Granin proteins, including chromogranin A (CHGA, CGA) and B (CHGB, CGB), provide a selective and condensing power for granule-targeted cargo molecules. Genetic and biochemical studies have shown that chromogranins (CGs) are fundamental regulators of granule biosynthesis (*27–29*). Intriguingly, ectopic expression of either CGA, CGB or other granin family members induces SG-like structures in non-secretory cells (*28, 30–32*). Experiments utilizing purified proteins or crude extracts of adrenal or pituitary glands showed that these molecules aggregate at acidic pH (5.5) and in the presence of millimolar concentrations of calcium (*33–35*). These observations led to the proposal of the “sorting by aggregation” or “sorting for entry” hypothesis in which aggregated granins and associated proteins exclude other soluble proteins in the TGN and promote SG biosynthesis (*27, 36–38*). However, the relevance of these observations in living cells is yet to be determined as the pH and calcium concentrations used in these studies are significantly beyond the physiological range (*39, 40*).

Liquid–liquid phase separation (LLPS) is increasingly recognized as a major principle of macromolecular organization within membrane-less compartments that enriches certain components and promotes specific biochemical reactions (*41–43*). The molecules that drive phase separation are referred to as scaffolds, whereas molecules that preferentially partition into the scaffolds are termed clients (*44–47*). The formation of these condensates is typically driven by multivalent interactions among proteins. One commonly identified element associated with phase separation in proteins is the intrinsically disordered region (IDR), an amino acid sequence characterized by low sequence complexity (*48, 49*). Simple charge patterns and the overall sequence composition of the IDRs are critical for phase separation. Other crucial factors controlling LLPS are the concentration and identity of macromolecules as well as the conditions of the microenvironment surrounding the condensates. Environmental conditions that influence the LLPS of soluble proteins include temperature, salt type and concentration, co-solutes, pH, and the volume excluded by others (*50*).

Here, we aimed to gain a comprehensive understanding of the mechanism underlying proinsulin sorting from the Golgi apparatus to ISGs of pancreatic β-cells as it remains an unresolved question in cell biology.

## Results

This study aimed to gain a comprehensive insight on mechanisms of proinsulin sorting at the TGN. We first analyzed insulin localization in a cell culture model of pancreatic β-cells (INS1 832/13) and observed insulin in punctate structures at the Golgi apparatus (Figure 1A). To confirm this result, we performed pulse-chase experiments with live cells expressing SNAP-tagged proinsulin (*29*) and monitored de novo synthesized proinsulin at the Golgi apparatus (Fig. 1B). SNAP-tagged proteins were observed in fluorescent structures similar to those observed by immunofluorescence microscopy (Fig. 1C). To determine whether these structures are located outside or inside the TGN, we reconstructed the volume of the TGN using 3D-rendering of confocal stacks from cells labeled using TGN38 and pan-insulin antibodies. Within the volume mask, punctate dots of insulin were observed, implying their presence in the TGN lumen (Figure 1D). We further confirmed the presence of electron-dense structures within the Golgi stack via transmission EM of mouse islets (Figure 1E). Proinsulin processing occurs during granule maturation (*51*), and processing enzymes must be co-sorted into vesicles targeting ISGs to facilitate proper insulin maturation. To test whether proteolytic enzymes colocalize with proinsulin puncta, we labeled the cells with pan-insulin and protein convertase 2 (PC2) antibodies; PC2 was found colocalized with the proinsulin puncta at the Golgi apparatus (Figure 1F).

**Figure 1:**
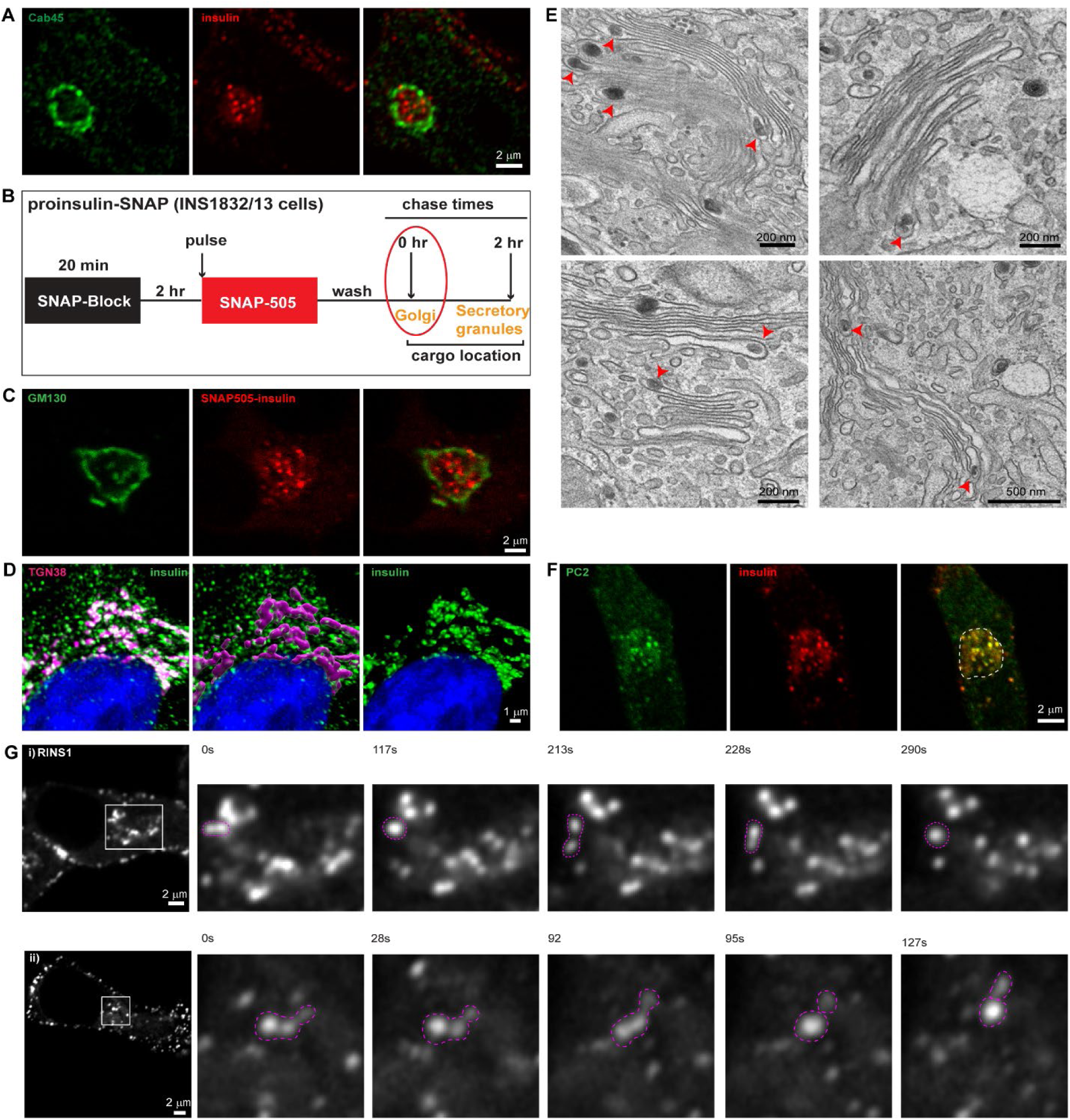
Insulin is distributed in dynamic punctate structures at the Golgi apparatus. **A)** Representative images from a single slice from a confocal image obtained from INS1 832/13 cells stained with the Golgi resident protein, Cab45 (green), which outlines the Golgi regions and insulin (pan-insulin antibody; red). Note the punctate structures of proinsulin seen in the perinuclear Golgi region. **B)** A schematic description of a pulse-chase assay in INS1 832/13 cells stably expressing SNAP-tagged proinsulin. Cells are initially incubated with a non-fluorescent blocking probe to mask the existing proteins in the cells. After 2 hours, cells are labeled with SNAP 505 and CLIP-TMR to mark the newly synthesized proteins (20 mins). After three washes in growth medium, cells are fixed immediately to monitor the pulse of new synthesized proinsulin arriving at the Golgi apparatus. **C)** Representative images from a single slice from a confocal image from INS1 832/13 cells expressing SNAP-tagged insulin (red) monitoring the insulin pulse arriving at the Golgi apparatus identified by the cis-Golgi marker GM130 (green). **D)** INS1 832/13 cells were immunostained for insulin (pan-insulin antibody; green), TGN38 (magenta) and counterstained with DAPI (blue). The middle image demonstrates 3D-rendering of the TGN38 volume mask which was used for image segmentation to specifically examine insulin staining within the TGN (right image). **E)** A panel of electron micrographs from mouse islets demonstrating condensed structures within the Golgi lumen**. F)** Representative images from a single slice from a confocal image of INS832/13 cells stained with the insulin processing enzyme protein convertase 2 (PC2; green) and insulin (pan-insulin antibody; red). Note the colocalization of proinsulin puncta with PC2 at the Golgi apparatus which is outlined in the merge image using dashed lines. **G)**A panel of images extracted from a movie from live imaging of INS832/13 cells transiently transfected with RINS1, a fluorescent insulin reporter construct. Dynamics of the punctate structures (pink dashed line) are captured in the image sequence where structures undergo fission or fusion events. Images shown here are single confocal slices upon imaging in the conventional confocal mode (i) or in the airy scan confocal mode (ii). Images have been smoothened using the function in ImageJ for visual representation purposes.

Incubation of cells with Brefeldin A (BFA) has been shown to result in disassembly of the Golgi apparatus and translocation of the associated proteins to the ER (*52*). Treatment of INS1 832/13 cells with BFA for resulted a dramatic change in the staining pattern of TGN38 (Supp Figure 1A), indicating disassembly of the TGN. Importantly, this was accompanied by disappearance of the perinuclear PC2 puncta implying these structures are associated with the Golgi apparatus. If the proinsulin/PC2 puncta are post-Golgi vesicles, they should remain in the perinuclear regions upon treatment with nocodazole, which generates Golgi ministacks (*53*). However, we found that PC2 was dispersed with the Golgi marker GM130 (Supp figure 1B), further suggesting the association with the Golgi apparatus.

Next, we investigated the dynamic nature of proinsulin puncta in the TGN. INS1 832/13 cells were transfected with RINS1, a dual fluorescently tagged form of proinsulin where mCherry is inserted within the C-peptide and super-folder GFP (sfGFP) is present at the C-terminus (*54*), and monitored using time-lapse live-cell imaging. The proinsulin puncta in the Golgi apparatus were highly dynamic and displayed fission and fusion events (Figure 1G). Overall, the observations reveal that proinsulin is organized in dynamic punctate structures at the Golgi apparatus.

### CGB and CGA undergo LLPS *in vitro*

Although the structure and dynamics of insulin puncta in the TGN displayed properties associated with liquid-like condensates, the polypeptide sequence of proinsulin did not show characteristics associated with proteins undergoing LLPS. Previous work has shown that CGs are significant regulators of proinsulin trafficking and that their depletion impairs insulin granule biosynthesis and glucose-stimulated insulin secretion (*29*). Analysis of the CGA or CGB sequences using the database PONDR (http://www.pondr.com) or AlphaFold (https://alphafold.ebi.ac.uk), respectively predicted that 50% of CGB and 90% of CGA are disordered (Figure 2A). We thus tested whether purified CGs form liquid-like condensates. To purify CGs, we generated CGA and CGB proteins containing sfGFP and His tags at their C-termini (using sfGFP with a monomerizing mutation; Supp figure 2A). Secreted proteins were purified from the culture medium of HEK293 cells stably expressing tagged proteins using Ni-NTA columns; proteins were obtained with high yields and purity (Figure 2C, D). To determine whether CGs form condensates, we changed the buffer from pH 7.4 to 6.1 to match the TGN lumen pH (*39*), and the protein solution was plated onto an imaging dish prior to analysis by fluorescence microscopy. We observed CGB condensates/droplets, whereas GFP remained soluble (Figure 2B, C). In contrast to CGB, purified CGA-GFP in solution did not form condensates when equilibrated at pH 6.1. However, condensate formation of CGA was induced by adding PEG 8000 to the protein solution (Figure 2D). Upon decreasing the protein concentration, the condensate sizes also decreased (Figure 2D).

**Figure 2:**
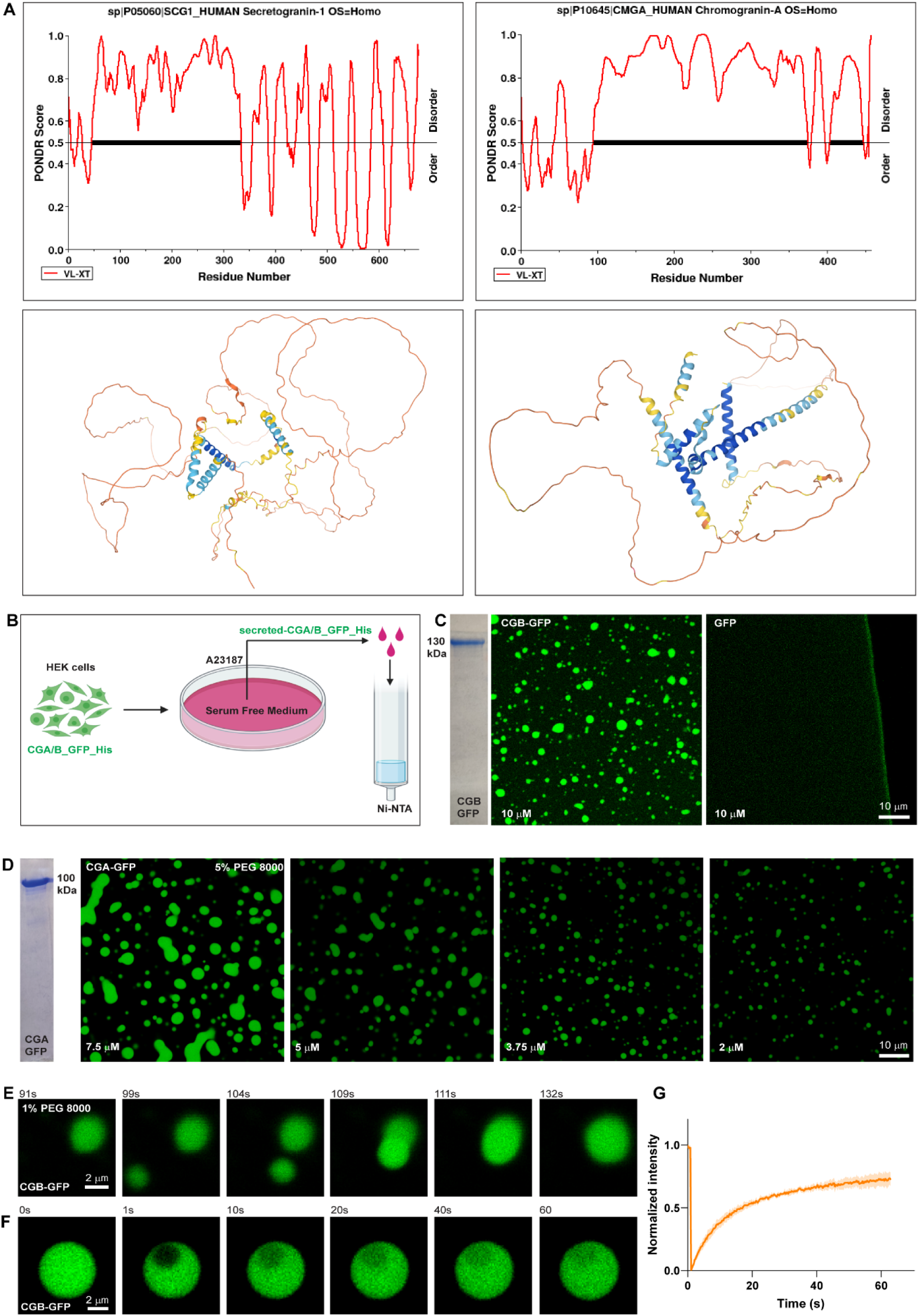
CGs undergo LLPS *in vitro*. **A)** Plots of CGB (left) and CGA (right) generated using PONDR depicting disordered regions in the proteins. While N-terminal half of CGB is completely disordered, almost 90 % of CGA shows disorder when analyzed using the VL-XT algorithm. Images below show the structural predictions of CGB and CGA using Alpha fold. **B)** Scheme used for purifying GFP and 6x-His tagged CGA or CGB, respectively. Stable lines expressing CGB-GFP and CGA-GFP under a doxycycline inducible promoter are treated with doxycycline and the calcium ionophore A23187 to induce secretion of the respective proteins in serum free medium, which is then used for purification using Ni-NTA affinity columns. **C)** Coomassie stained gel depicting purified CGB-GFP. The images show droplets of CGB-GFP (left) at 10 µM protein concentration at pH 6.1 without crowding agent. The same concentrations of super folder-GFP does not form droplets at pH 6.1 (right). **D)** Coomassie stained gel depicting purified CGA-GFP. Images showing different droplets of CGA-GFP at different protein concentrations at pH 6.1 in presence of 5% PEG 8000. Note that the size of the condensates decreases with decreasing protein concentration. **E)** A panel of images extracted from a movie showing the biophysical behavior of CGB-GFP (2.5 µM) droplets induced by 1% PEG8000. Two droplets, which come in proximity undergo fusion and the larger droplet subsequently relaxes into a spherical shape. **F)** A panel of images monitoring recovery of fluorescence of CGB-GFP droplets after bleaching a small region within the droplets. Note the rapid recovery of fluorescence (more than 60%) in the bleached region within a minute. **G)** Graph quantifying the fluorescence recovery in time. Data represented as mean +/- s.d. (error blanket) from eight independent droplets.

Live-cell imaging of CGB-GFP droplets to determine whether they exhibit liquid-like properties showed that two or more condensates in proximity display fusion and relaxation to a spherical shape (Figure 2E). Moreover, when a small region within the CGB condensate was photobleached, more than 50% of the fluorescence recovered within one minute, indicating high mobility of proteins within the condensates (Figure 2F, G). CGA condensates induced with PEG 8000 displayed similar liquid-like behavior (Supp figure 2B, C). These data demonstrate that purified CGs undergo LLPS *in vitro* and that CGB has a higher potential for phase separation than does CGA.

### The TGN pH drives CG LLPS while divalent cations are dispensable

Secretory proteins are exposed to changes in the chemical environment as they travel through the sub-compartments (milieu) of the biosynthetic pathway. Having established a straightforward microscopy-based *in vitro* assay to monitor CG condensation, we investigated the impact of pH on this process by using the intrinsic pH difference between the ER and TGN compartments. We analyzed droplet formation by fluorescence microscopy using purified GFP-tagged CGs with buffer conditions of pH 7.3 (ER milieu) or 6.1 (TGN milieu) (*39*). Interestingly, CGA (with crowding reagent) and CGB formed droplets at pH 6.1 but not 7.3. This demonstrates that CG condensates are formed at a pH consistent with the TGN milieu, suggesting a physiological triggering mechanism for CG condensation.

Given that CGs aggregate in the presence of high calcium concentrations (*34–36*), we tested the influence of calcium on LLPS by incubating GFP-tagged recombinant proteins at pH 6.1 with varying concentrations of calcium and analyzing droplet formation. CGB-GFP (10 µM) solution, centrifuged to pre-clear existing droplets, was diluted to 2.5 µM in the presence of varying concentrations of calcium. The steady-state calcium concentration in the TGN is estimated to be 130 µM (*40*); surprisingly, we did not observe droplet formation for either CGA or CGB at 250 µM calcium (Supp figure 3A, C), which is closer to the physiological concentrations in the TGN than are the millimolar-range concentrations used in previous studies. Instead, calcium (5 mM) induced the formation of CGB condensates, which was significantly potentiated at 20 mM calcium based on the size and density of droplets (Figure 3B; Supp Figure 3A, B). Although CGB binds to calcium (*55*); no structural binding motif, such as an EF-hand domain, has been identified. Consequently, we questioned whether such condensation under high calcium concentrations is specific or can be induced by other divalent cations. Magnesium and manganese at 20 mM induced LLPS of CGB (Figure 3B). However, at 5 mM, magnesium failed to enhance CGB LLPS, whereas manganese induced an increase in CGB condensate size (Supp figure 3B). Since CGA does not form condensates on its own in solution, we added calcium to determine if it results in LLPS of CGA without crowding agents. However, even at high concentrations (40 mM), we did not detect CGA droplet formation at pH 6.1 (Supp Figure 3C).

**Figure 3:**
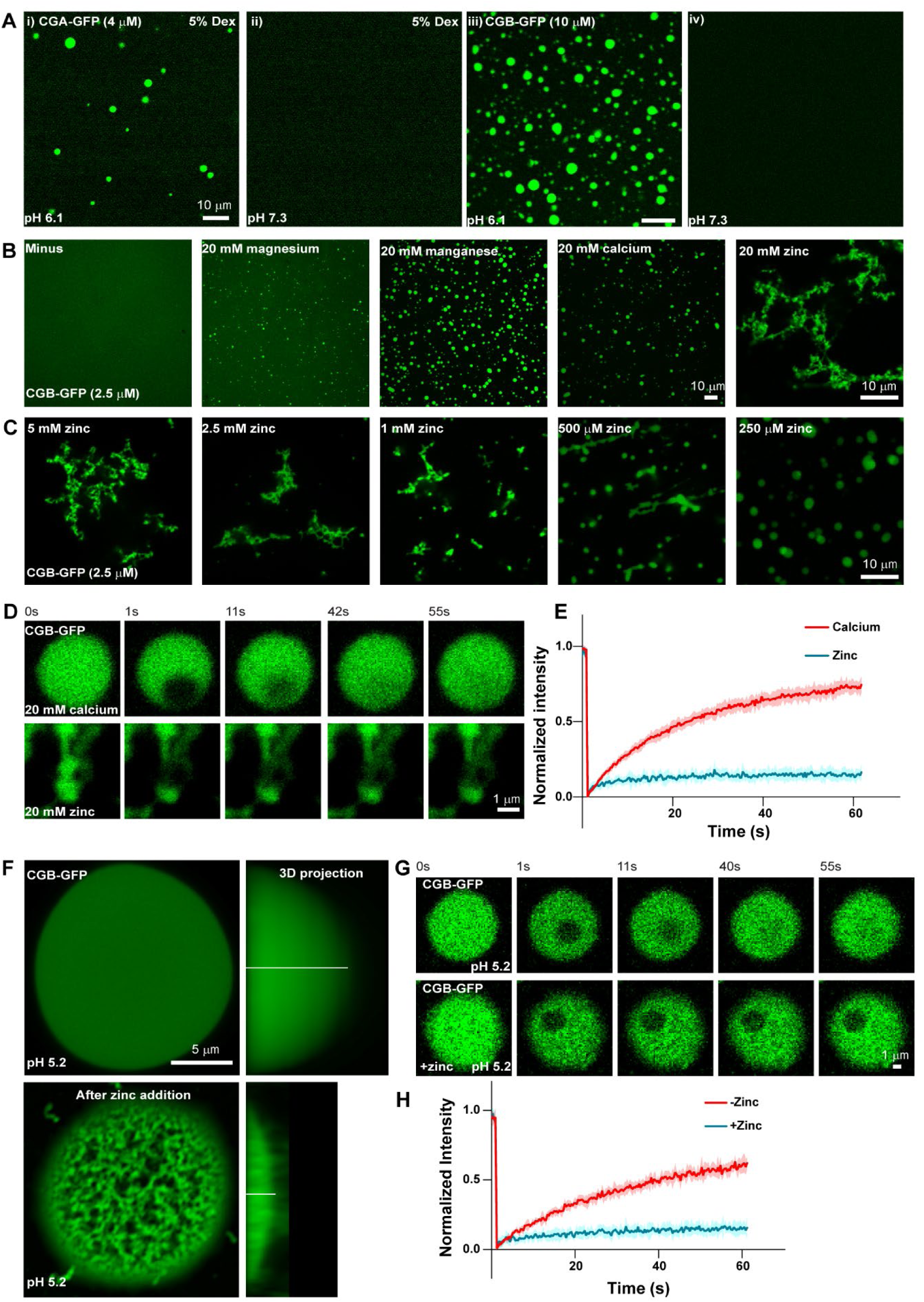
Factors influencing LLPS of CGs. **A)** Representative images of solutions containing CGA-GFP (i, ii) or CGB-GFP (iii; iv) buffered at either pH 6.1 (I, iii) or pH 7.3 (ii, iv). Droplet formation occurs at pH 6.1 and not at pH 7.3. Droplets of CGA-GFP were induced at 4 µM protein concentration in presence of 5% dextran, while 10 µM CGB-GFP formed droplets without crowding agents at pH 6.1. **B)** Representative images of CGB-GFP (2.5 µM) protein without any divalent cation (minus) or in presence of 20 mM magnesium, manganese, calcium, and zinc respectively. CGB-GFP solution (approximately 10 µM concentration) was centrifuged to preclear of existing droplets and diluted to a final concentration of 2.5 µM. While, magnesium, manganese and calcium induce CGB-GFP droplets, zinc induces formation of insoluble aggregates. In terms of droplet formation potential manganese > calcium > magnesium. **C)** Representative images of CGB-GFP (2.5 µM) in presence of different concentrations of zinc. At high concentrations zinc induces formation of insoluble aggregates but at low concentration it induces CGB-GFP droplets. **D, E)** Snapshots captured at different time points after photobleaching a region within a calcium induced CGB-GFP droplet (top panel) or a zinc induced CGB-GFP aggregate are shown in D. While there is more than 60% fluorescence recovery within the bleached region in the calcium induced droplet, recovery within the bleached region in the aggregate is only 10% as seen in the graph in E. Red curve denotes recovery of calcium induced droplets and blue curve denotes zinc induced aggregates. Data is represented as mean +/- s.d. (error blanket) from at least six droplets/aggregates. **F)** Images on the left side show a single plane from an airy-scan confocal image of the CGB-GFP condensate at pH 5.2 <u>before</u> (top panel) and after addition of 20 mM zinc (bottom panel). Zinc induces aggregation within a preformed droplet and is accompanied by shrinkage in volume as seen from the 3D-projections on the right. **G, H)** A panel of images from a FRAP experiment involving bleaching of a small region within CGB-GFP droplets formed at pH 5.2 (top panel) and after treating them with zinc (bottom panel). Graph in H shows the comparison of FRAP recovery curves of CGB-GFP droplets at pH 5.2 with (blue curve) and prior to zinc addition (red curve). While the fluorescence recovery in the bleached region within CGB-GFP droplets without zinc addition is approximately 50%, droplets get hardened after zinc addition and recovery is approximately 15%. Data is represented as mean +/- s.d. (error blanket) from at least seven droplets in each condition.

Zinc is essential for hexamerization of mature insulin (*56*) and is present at high concentrations (mM level) in SGs due to the localization of zinc transporters (*21, 57, 58*). To test its impact on droplet formation, fluorescent proteins were incubated with high concentrations of zinc (20 mM), after which we observed filamentous aggregates of CGB (Figure 3B). Partial fluorescence recovery after photobleaching (FRAP)—bleaching a small region in the zinc-induced aggregates or calcium-induced condensates to monitor fluorescence recovery within the bleached spot—was used to confirm whether they are indeed insoluble solid aggregates. While calcium-induced condensates recovered fluorescence in the bleached region by more than 50% within a minute, there was almost no recovery of photobleached zinc aggregates (Figure 3D, E). To further investigate the effects of zinc on CGB, the zinc concentration was titrated. We observed a mixture of aggregates and condensates at 1 mM zinc and droplets below 1 mM zinc (Figure 3C). In pancreatic β-cells, as the immature SGs mature, there is a further drop in their luminal pH to approximately 5.2 (*39*), which is more acidic than that of the TGN lumen. At pH 5.2, CGB continued to form condensates; however, if the pre-formed condensates were treated with 20 mM zinc, an immediate hardening of the condensates accompanied by massive volume shrinkage occurred (Figure 3F). FRAP measurements confirmed differences in condensate fluidity before and after the addition of zinc (Figure 3G, H).

The above data reveal that an acidic pH of 6.1 is the primary driver for CG LLPS and that the presence of divalent cations modulates the LLPS of CGB, contradicting the current understanding that calcium is required for inducing CG aggregation in the TGN. Calcium and other divalent cations can boost the inherent ability of CGB to undergo LLPS, but they are not necessary for the reaction.

### Proinsulin and CGB are co-sorted into TGN-derived transport carriers

Based on the CGB condensation results we hypothesized that it is a necessary step for the sorting of proinsulin at the TGN. Depletion of CGB in INS1 832/13 cells results in impaired proinsulin export from the TGN (*29*). A similar result was obtained in islets derived from CGB knockout mice (*59*). If our hypothesis that CGB condensation sorts proinsulin in the TGN is correct, both proteins should colocalize and bud into the same transport carrier exiting the TGN. To address this, we established a pulse-chase approach to monitor the simultaneous export of proinsulin and CGB from the TGN in living cells. We expressed CLIP-tagged CGB in INS1 832/13 cells stably expressing SNAP-tagged proinsulin (Figure 4A). Newly synthesized SNAP-proinsulin and CLIP-CGB entering the TGN were labeled with their respective fluorescent analogs (t = 0) and observed to colocalize (Fig. 4A). Following a 2-hour chase of the respective fluorescent SNAP or CLIP probes, both proteins colocalized in cytoplasmic granules, indicating that they are sorted into the same transport carriers exiting the TGN (Figure 4A). Following a 2-hour chase of the respective fluorescent SNAP or CLIP probes, both proteins colocalized in cytoplasmic granules, indicating they were sorted into the same transport carriers exiting the TGN (Figure 4A). Moreover, proinsulin at steady-state showed colocalization with CGB, CGA, and secretogranin III in the TGN, TGN-derived vesicles, and cytoplasmic granules (Figure 4B-D).

**Figure 4:**
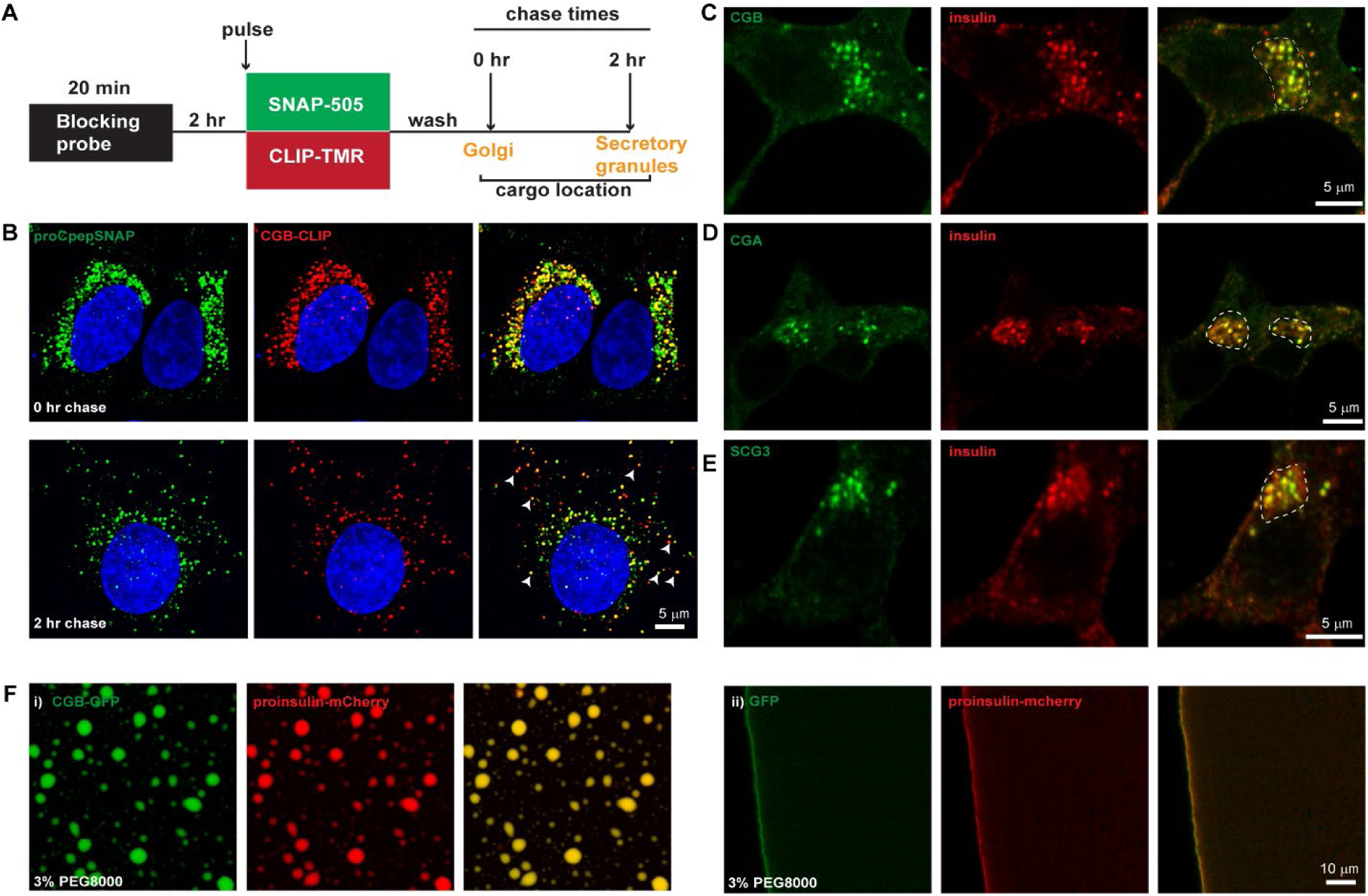
Proinsulin co-traffics with CGB *in vivo* and is recruited to droplets *in vitro*. **A)** Schematic depiction of dual pulse chase experiment in INS1 832/13 cells expressing SNAP tagged insulin (proCpepSNAP; SNAP tag and CLIP tagged CGB. Cells are initially incubated with a non-fluorescent blocking probe to mask the existing proteins in the cells. After 2 hours, cells are incubated with medium containing with SNAP 505 and CLIP-TMR to label the newly synthesized proteins (20 mins). After three washes in growth medium, cells are fixed immediately (0 h chase) when majority of the cargo is at the Golgi apparatus or after a chase of 2 hours where most of the cargo has moved to the SG in the cytoplasm. **B)** Top panel shows confocal images form INS832/13 cells expressing SNAP tagged insulin (proCpepSNAP; green) and CLIP tagged CGB in red and fixed immediately after labeling with fluorescent probes, SNAP-505 and CLIP-TMR to monitor the Golgi resident (peri-nuclear) pool of the proteins. Bottom panels show images after a 2-hour chase and the arrows point to some of the colocalizing structures which are cytoplasmic SGs. **C-E)** INS832/13 cells fixed and labeled with antibodies to insulin in or CGB (A), CGB (B) and SCG3 (C) in green. Insulin puncta at the Golgi apparatus in the perinuclear region, outlined using the dashed lines in the merge colocalize with each of the granin proteins. For CGB and CGA, images are a single slice from a confocal stack and in case of SCG3, an average projection from three consecutive slices from a confocal image. **F)** CGB-GFP (2.5 µM; green) was mixed with mcherry tagged proinsulin (1 µM; red) in (i) and droplets were induced at pH 6.1 using 3% PEG 8000. Tagged proinsulin gets recruited to the CGB-GFP droplets as evident from the colocalization image. When GFP (2.5 µM; green) is mixed with mCherry tagged proinsulin (1 µM; red) in (ii), no droplets are seen either with GFP or proinsulin even after addition of 3% PEG8000, indicating that GFP or mCherry-proinsulin are incapable of forming droplets on their own at these concentrations.

To gain further insights into the relationship between proinsulin and CGB, we applied our well-established *in vitro* system mimicking TGN conditions. CGB-GFP (2.5 µM final concentration) and mCherry-tagged proinsulin (1 µM final concentration) were combined at pH 6.1 and 3% PEG 8000 was used to induce robust CGB condensate formation. Recruitment of proinsulin to the CGB condensates was readily observed (Figure 4F); no co-condensation of GFP and proinsulin occurred (Figure 4F). Based on these findings and previously published results, we propose that CGs co-condense in the luminal milieu of the TGN. Moreover, CGB condensation results in proinsulin co-segregation and export from the TGN.

### Ectopic expression of constitutively secreted proteins results in their targeting to SGs and co-secretion with insulin

Our findings so far suggest that CG condensates containing proinsulin and processing enzymes bud out vesicles from the TGN to form immature secretory granules, which in turn mature into fully functional insulin granules. By necessity, molecules destined for lysosomes or constitutively secreted proteins would be segregated from the CG condensates in the TGN. To monitor the co-segregation or segregation of cargo molecules within the TGN lumen, we examined a subset of proteins destined for locations other than SGs. Significant colocalization of proinsulin and carboxypeptidase E, an SG cargo and proinsulin processing enzyme, was observed (Figure 5A, D). On the other hand, comparison of the localization of proinsulin and the cargo receptors of lysosomal hydrolases (M6PR or IGF2R) revealed that these proteins segregate within the TGN lumen (Figure 5B, C and Figure 5E, F). Similarly, endogenous cathepsin B was segregated from proinsulin in the TGN (Supp figure 4B). This suggests that lysosomal hydrolases and their receptors exit from different regions of the TGN.

**Figure 5:**
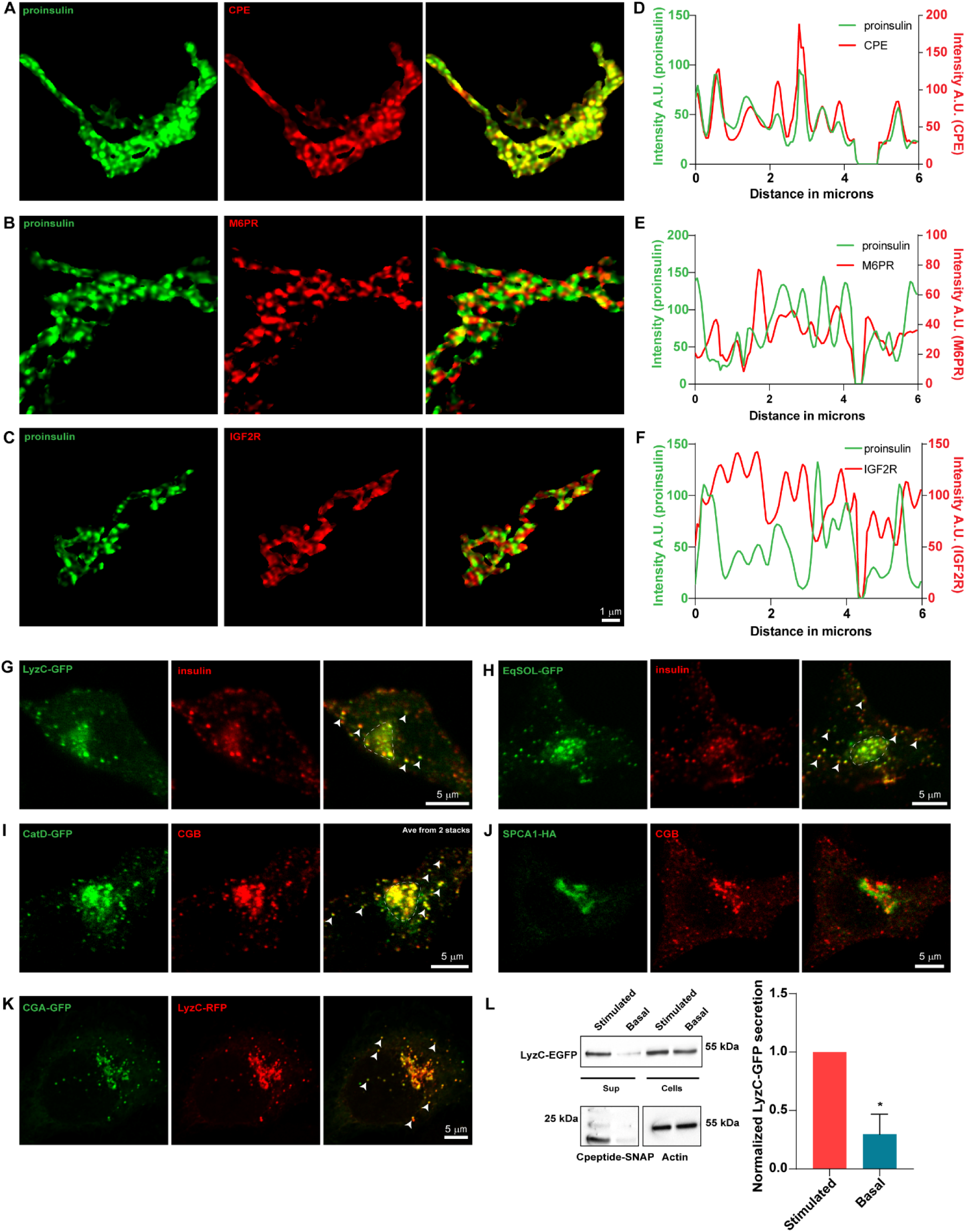
Ectopic expression of soluble secreted proteins in INS832/13 cells results in their routing to insulin granules. **A-F)** INS1 832/13 cells were immunostained for insulin (pan-insulin antibody; green), TGN38 (shown in Supp Figure 4A), CPE (**A**, **D**), M6PR (**B**, **E**), or IGF2R (**C**, **F**). (**A**-**C**) Image segmentation of the TGN38 volume was used to identify Golgi-staining regions. (**D**-**F**) Line intensity profiles of the Golgi staining regions for each protein are shown. **G, H)** Representative images from INS832/13 cells expressing LyzC-GFP (E; green) or EqSOL-GFP (F; green) and stained with insulin antibody (red) to observe the localization of the ectopically expressed proteins with respect to insulin granules. Images are average projections from two slices from a confocal stack. Arrowheads point to cytoplasmic insulin granules which also shows the presence of LyzC-GFP and EqSOL-GFP respectively, in E and F. **I)** Representative images from INS1 832/13 cells stably expressing CatD-GFP (green) and labeled with CGB antibody to observe the localization of ectopically expressed CatD-GFP with respect to SGs. Images are average projections from two slices from a confocal stack. Arrowheads point to some of the cytoplasmic SG, which shows colocalization of CGB and CatD-GFP. **J)** Representative images from INS832/13 expressing HA-tagged version of the calcium ATPase, SPCA1 (green) and stained using CGB antibody (red). Images are a single slice from a confocal stack. Note that overexpressed SPCA1 remains localized at the Golgi apparatus with no signal seen from the CGB containing SGs. **K)** Representative images from HeLa cells stably expressing CGA-GFP and transfected with LyzC-RFP. Images are a single slice from a confocal stack imaged in the airy-scan mode. The arrowheads point to some of the ectopic granule-like structures seen in HeLa cells upon expression of CGA-GFP. Note that LyzC-RFP gets routed to these ectopic granule-like structures. **L)** Western blot at the top shows bands for LyzC-GFP, probed using α-GFP antibody, in supernatant and lysates from INS1 832/13 cells stable expressing SNAP-tagged proinsulin. The basal condition represents cells grown in 3 mM glucose in serum free medium and the stimulated condition represents cells grown in 15 mM glucose in serum free medium, also containing 35 mM potassium chloride. Note the stronger band intensity in the supernatant in stimulated condition compared to the basal condition, although the levels in cell lysates are the same. The blot in the bottom left detects the presence of SNAP tagged C-peptide, probed using α-SNAP antibody, which is used as a proxy to measure insulin secretion. Again, the signal intensity of the band is stronger in stimulated condition as compared to the basal condition. The blot on the bottom right depicts actin bands in cell lysates obtained from basal and stimulated conditions. The graph quantifies secretion of LyzC-GFP normalized with levels in cell lysates in basal and stimulated conditions. Data is represented as mean +/- s.d. from three independent experiments. *\* p< 0.05*.

Next, we wondered whether constitutively secreted proteins segregate from proinsulin CGA droplets, which would align with the “sorting for entry” model (*38*). Cab45 sorts a subset of proteins, including LyzC, to the cell surface in HeLa cells (*60*). Therefore, proinsulin localization was compared with that of ectopically expressed LyzC or EqSOL, a soluble bulk flow marker. Unexpectedly, we observed colocalization of these proteins with proinsulin, and both LyzC and EqSOL were transported to SGs in insulin-secreting cells (Figure 5G, H). We then analyzed localization of the ectopically expressed lysosomal hydrolase cathepsin D (CatD) in INS1 832/13 cells. In contrast to the endogenous hydrolase, ectopically expressed CatD-sfGFP was not targeted to the lysosome but instead routed to insulin granules (Figure 5I). However, this effect appears to be limited to soluble TGN cargoes, as ectopic expression of HA-tagged SPCA1, a transmembrane protein localized to the TGN, did not result in SPCA1 and proinsulin colocalization but instead remained TGN localized (Figure 5J). To further investigate whether routing to SGs depends on CGs, we expressed mCherry-tagged LyzC and CGA-sfGFP in HeLa cells, which are naturally deficient in CGs and SGs. We detected ectopic secretory granules in HeLa cells expressing CGA-sfGFP and observed the routing of LyzC-mCherry to these granules (Figure 5K). These data indicate that CG condensates can non-specifically recruit client proteins and that CG expression overrides existing pathways for sorting soluble proteins to favor SG accumulation.

Since overexpression of secreted soluble proteins leads to their routing to the SGs in INS1 832/13 cells, we assessed whether they are also secreted in response to stimulus, as is the case with insulin (*61*). Indeed, at basal levels (3 mM glucose), we observed a minimal amount of basal secretion in cells expressing the constitutively secreted protein LyzC-GFP and SNAP-tagged C-peptide (proxy for insulin secretion). However, upon glucose (15 mM) and potassium chloride (35 mM) stimulation of INS1 832/13 cells, both LyzC-GFP and insulin were co-secreted (Figure 5L). These results suggest that CG condensates recruit soluble proteins by nonspecific mechanisms that do not rely on the sequence or structural motif of the client. The co-condensation of soluble proteins with CGs results in their routing to SGs, which in turn allows them to be co-secreted with insulin in response to glucose stimulation.

### LyzC gets recruited to granin condensates *in vitro*

We then determined whether other soluble proteins such as LyzC and CatD are recruited to CGB condensates *in vitro* and whether the liquid-like droplets or aggregates of CGB are equally competent to recruit cargo. CGB and Cy3-tagged LyzC were combined at 2.5:1 molar ratios and droplet formation was induced using calcium. Under these conditions, Cy3-LyzC was recruited to the CGB condensates (Figure 6A, B). Since we observed CGB aggregates only in the presence of zinc, we tested whether zinc-induced CGB aggregates recruit Cy3-LyzC. In contrast to the current view that aggregated CGs recruit clients, we did not detect Cy3-LyzC recruitment into the zinc-containing CGB aggregates (Figure 6A, B). In a similar experiment, we confirmed that Cy3-tagged CatD was also recruited to the CGB condensate, supporting a client recruitment mechanism that is independent of a specific sequence (Figure 6E).

**Figure 6:**
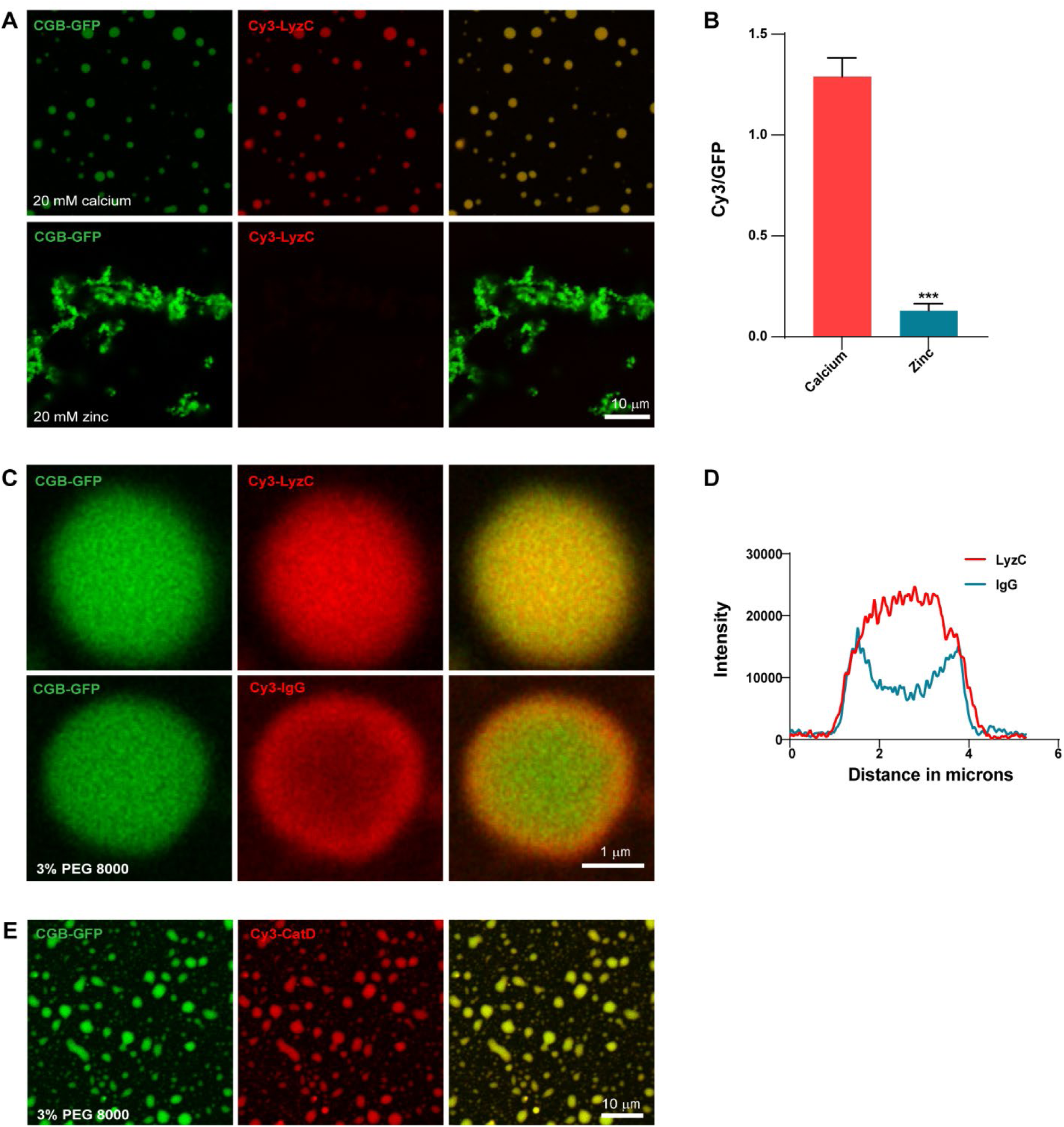
Client recruitment *in vitro*. **A, B)** CGB-GFP (2.5 µM; green) was mixed with Cy3-tagged LyzC (1 µM; red) and was subsequently followed by addition of 20 mM calcium to induce liquid like droplets or with 20 mM zinc to induce aggregates. While calcium induced CGB-GFP droplets recruit Cy3-LyzC, the zinc induced aggregates fail to recruit it. Bar graph is B quantifies the ratio of intensities in Cy3 v/s GFP channel, imaged at same acquisition settings, to monitor recruitment of LyzC in relation to amount of CGB in the droplet (red) v/s aggregate (blue). Data is represented as mean +/- s.d. from at least thirty droplets or regions within an aggregate. *^***^ p< 0.001*. **C, D)** CGB-GFP (2.5 µM; green) was mixed with Cy3-tagged LyzC (1 µM; red; top panel) or with Cy3-tagged Igg (1 µM; red; bottom panel) and droplet formation was induced using 3% PEG8000 to monitor the recruitment of these cargoes into the CGB droplets. While LyzC, which is 14.3 kDa in size shows a uniform recruitment within the CGB droplet, 150 kDa IgG, only shows recruitment at the edges of the CGB droplets. The images are a single slice of a confocal image to image through the cross section of a droplet from the center and were smoothened for representation purposes using the function in Image J. Graph in D, shows line scans in the Cy3 channel again taken approximately from the center of the droplet along the longest axis. For Cy3-LyzC, the intensity continues to rise as we move from the edge to the center with plateauing in the center of the droplet, while for Cy3-Igg there is a sharp decline in intensity in the center of the droplet, consistent with their recruitment profile. **E)** CGB-GFP (2.5 µM; green) was mixed with Cy3-tagged CatD (1 µM; red) and droplet formation was induced using 3% PEG8000 to monitor the recruitment of clients into the condensates. Note that Cy3-CatD shows a strong recruitment to the droplets.

Next, we investigated how CG condensates select cargo molecules. LyzC is a small protein (14 kDa) and the molecular weight of CatD is approximately 45 kDa. We thus assessed whether the small size of the protein allows entry into the CGB condensates by comparing Cy3-LyzC and Cy3-labeled rabbit IgG (MW 150 kDa) recruitment. Based on the intensity profile, LyzC filled the complete volume of the CGB condensate, whereas IgG only populated the edges of the CGB condensates, with little signal detected in the center of the droplet (Figure 6C, D). This suggests that CGB condensates enrich smaller molecules and exclude those that have a high molecular weight.

These results suggest that while CGB aggregates cannot recruit cargo and must be in a liquid-like state to do so, CGB condensates can recruit small sized clients independent of any sequence or structural features.

### Truncation of CGB impacts phase separation potential and the ability to generate ectopic granules

While we have just begun to understand the molecular mechanisms governing LLPS, there is no general defining factor that drives LLPS. To narrow down the region in CGB driving LLPS, we truncated the CGB protein into distinct parts based on its predicted structure (PONDR database); the N-terminal domain of CGB (amino acids: 46–334 aa) is highly unstructured (high degree of disorder) compared with the C-terminal (amino acids: 335-667; Figure 7A). Hence, sfGFP- or mCherry-tagged truncation mutants of CGB, CGB_N term-GFP and CGB_C term-mCherry, were generated. For each mutant, the signal sequence of CGB was incorporated at the N-terminus to allow transit through the secretory pathway. Neither of the truncation mutants displayed LLPS at pH 6.1 (Supp figure 5A); however, upon addition of 3% PEG 8000, the N-terminal was observed to undergo a significantly more potent LLPS than that of the C-terminal (Figure 7B). There was a difference in area covered by the droplets observed for each mutant, with the effects being much stronger for the CGB C-terminal fusion protein. (Figure 7C, E). The inability of the CGB C-terminal to undergo LLPS was not due to the mCherry tag because the sf-GFP-tagged version displayed the same phenotype (Supp figure 5B). Interestingly, when the N- and C-terminal truncation mutants were combined and phase separation was induced in the presence of 3% PEG 8000, the mixture displayed phase separation comparable to that of full-length CGB-GFP (Figure 7D, E). These results suggest that while most of the information responsible for mediating LLPS of CGB is encoded by the N-terminal domain, the C-terminal is required for optimal function.

**Figure 7:**
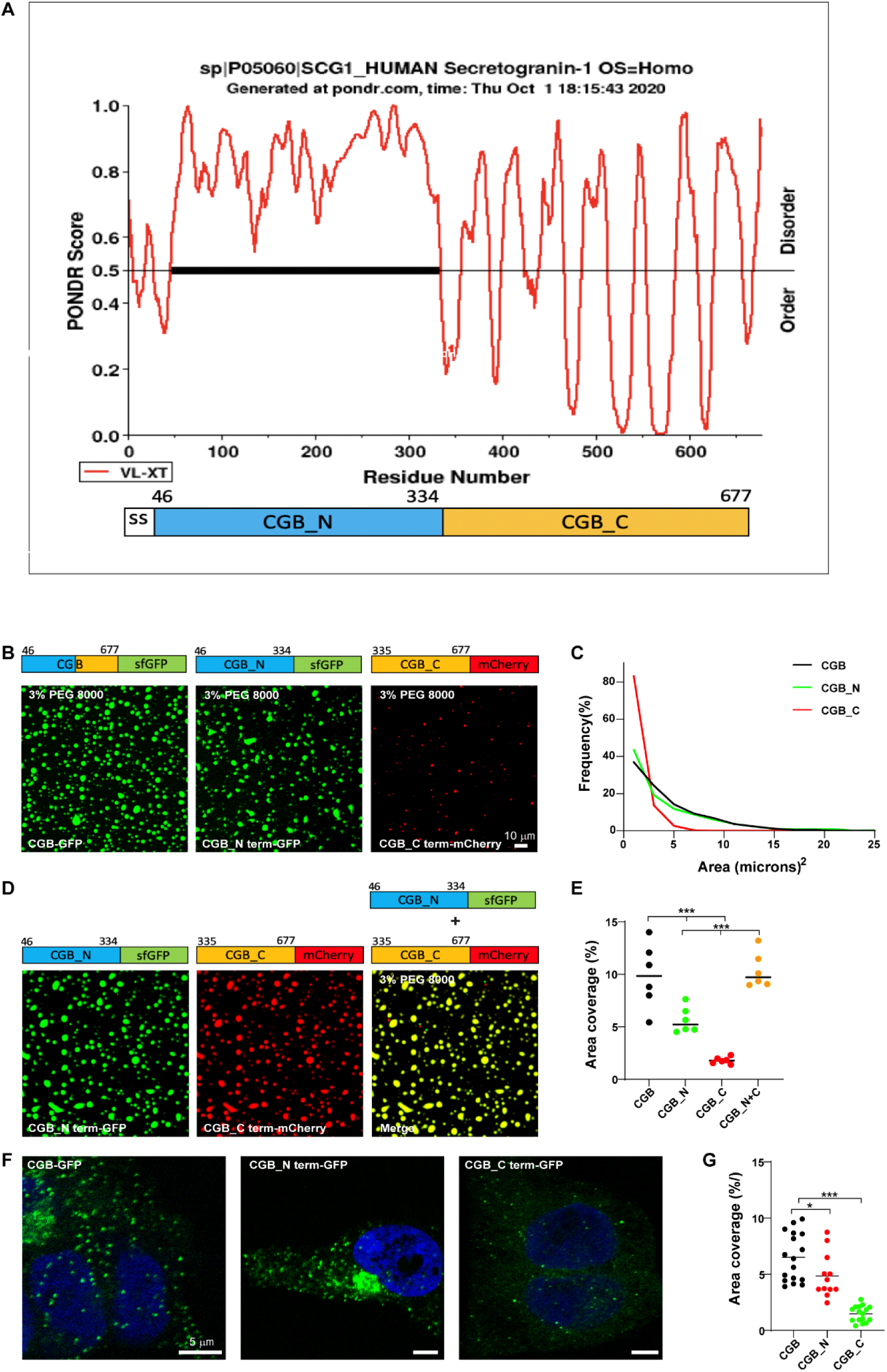
*In vitro* and *in vivo* phenotypes associated with truncation mutants of CGB. **A)** Plots of CGB generated using PONDR (VL-XT algorithm) depicting disordered regions in the protein. Based on the PONDR scores, amino acids 46-334 are highly unstructured, thus showing a high degree of disorder compared to residues 335-677. **B)** Representative images showing a comparison of condensates of full length CGB-GFP, CGB_N term-GFP and CGB_C term-mCherry respectively. Droplet formation was initiated at pH 6.1 with 2 µM protein in presence of 3% PEG8000. **C)** Histogram represents the frequency distribution of size of condensates for each of the three proteins. Area was quantified from at least one thousand droplets for each condition. Note that the size of the droplets is larger for CGB-GFP, CGB_N term-GFP compared to CGB_C term-mCherry. **D)** Representative images of droplet formation obtained upon mixing CGB_N term-GFP (green) and CGB_C term-mCherry (red). Droplet formation was initiated at pH 6.1 upon mixing 2 µM protein of each protein in presence of 3% PEG8000. CGB_N term-GFP and CGB_C term-mCherry cooperatively form larger droplets together. **E)** Graph quantifies area occupied by all the droplets in a field of view from a microscopy image. Data was pooled from 6 field of views. CGB-GFP > CGB_N term-GFP > CGB_C term-mCherry. Upon mixing CGB_N term-GFP with CGB_C term-mCherry, there is a restoration in the area covered by the condensates to the levels of the full-length protein. *^***^ p< 0.001*. **F)** Representative images of HEK293 cells expressing CGB-GFP (left), CGB_N term-GFP (middle) and CGB_C term-GFP (right) and induced with doxycycline for ten hours. Images were taken on a confocal microscope to observe ectopic granule-like structures in HEK293 cells, and the bottom-most plane of the cells plated on the coverslips was imaged. While CGB-GFP and CGB_N term-GFP expression induces formation of ectopic granules in HEK293 cells, much lesser granules are seen upon expression of CGB_C term-GFP. **G)** Graph quantifies the percentage area occupied by the ectopic granules as a fraction of total cell area. Data was obtained from at least 12 cells pooled from two independent experiments. *\* p< 0.05, ^***^ p< 0.001*.

To determine the contributions of LLPS to SG biogenesis, we took advantage of the CG protein characteristic of inducing ectopic granule-like structures when expressed in non-secretory cells (*30, 62*). For this purpose, we used stable HEK293 cell lines expressing CGB-GFP, CGB_N term-GFP, and CGB_C term-GFP and induced expression using doxycycline. We observed that ectopic expression of both full-length and N-terminal CGB induced biogenesis of ectopic granules, whereas the C-terminal CGB did not (Figure 7F, G). Altogether, these results reveal that the potential of CGB to undergo LLPS is directly linked to its ability to induce SG formation. Thus, CG LLPS is not only critical for sorting soluble proteins within the TGN lumen but also for driving SG biogenesis.

### Electrostatic interactions mediate LLPS of CGB

Electrostatic interactions have been shown to drive LLPS (*63*). In the case of the protein Ddx4, the arrangement of its amino acids into clusters of similarly charged residues has been shown to drive LLPS (*64*). While CGB is rich in acidic amino acid residues, a striking feature in the CGB protein are five clusters of negatively charged amino acids (Figure 8A). There are also dibasic sites that have also been shown to be proteolytic cleavage sites, with most present within the C-terminus. Based on these features, we hypothesized that electrostatic interactions drive the LLPS of CGB.

**Figure 8:**
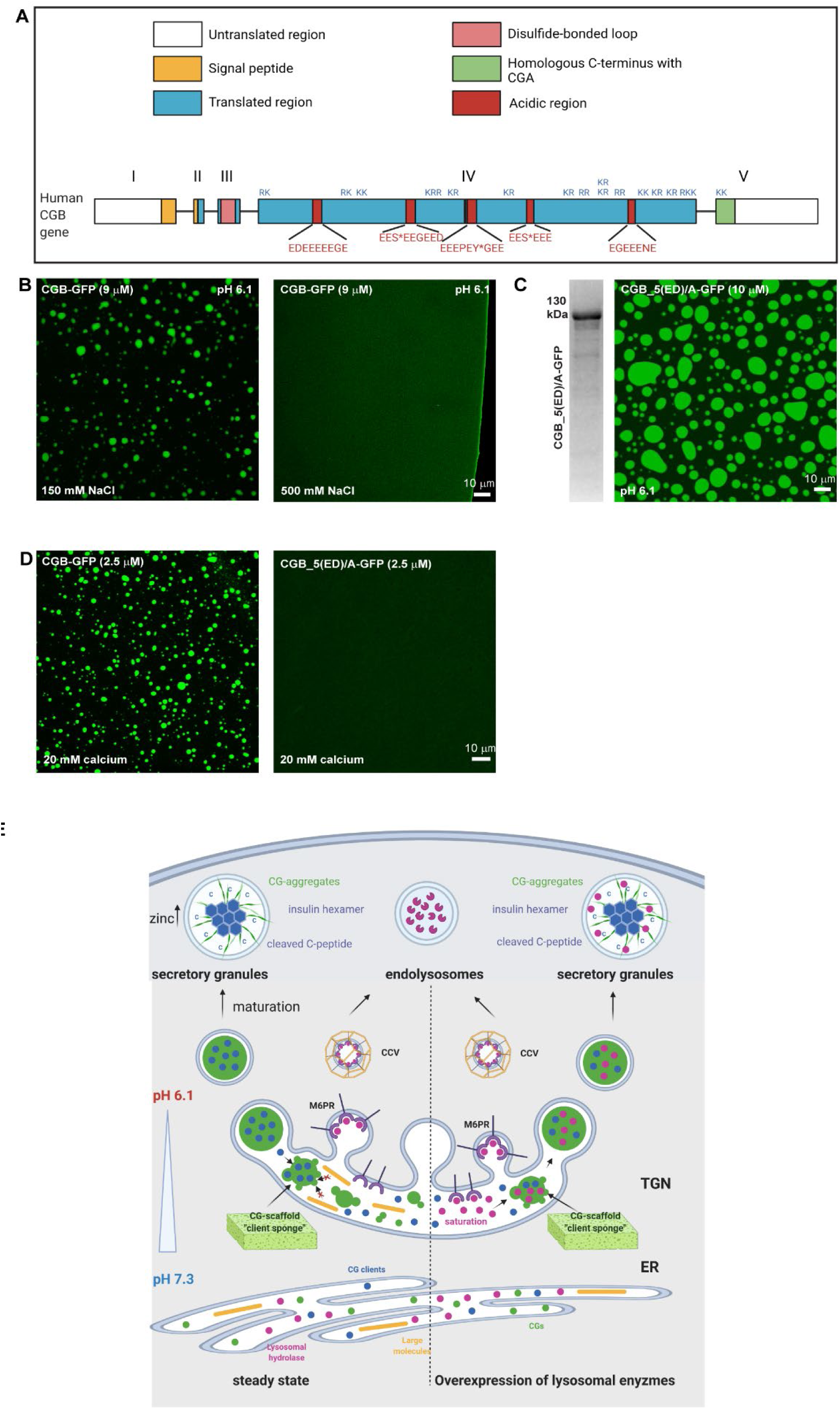
Effects of salt concentration and acidic amino acids on LLPS of CGB. **A)** Schematic representation of CGB, depicting 5 exons. Patches in red are stretches enriched in acidic amino acids. S* and Y* denote phosphorylation and sulfation respectively on these residues. KR or RK are the dibasic sites in the protein which are predominantly concentrated in the C-terminal half of the protein. **B)** Representative images of CGB-GFP equilibrated at pH 6.1 in presence of either 150 mM NaCl (physiological salt; left) or 500 mM NaCl (high salt; right). Droplet formation is only seen in presence of physiological salt concentration of 150 mM, but not seen at high (500 mM) NaCl concentrations. **C)** Representative Coomassie stained image of mutant form of CGB, CGB_5(ED)/A-GFP to depict its purity. In this mutant the 5 (ED) stretches have been replaced to alanine. The fluorescence image shows condensates of CGB_5(ED)/A when equilibrated at pH 6.1 at 10 µM protein concentration. **D)** Comparison of CGB-GFP and CGB_5(ED)/A-GFP in presence of 20 mM calcium concentrations. Note that at 2.5 µM protein concentration, calcium can induce droplet formation with only CGB-GFP, but now with CGB_5(ED)/A-GFP. **E)** Model presented summarizes our findings on the role of LLPS in receptor independent cargo delivery from the TGN to the SGs. CGs undergo LLPS in the milieu of the TGN, behaving like a “cargo sponge” and recruit clients like proinsulin by the virtue of their relative abundance to the condensates. Larger molecules which can only attach the condensate from edges would not be enriched in the CG condensates. Lysosomal hyrolases would be sorted to the endolysosomes via the mannose-6-phosphate receptor pathway. However, upon overexpression, the receptors are saturated and hence owing to their smaller size and high abundance at the TGN, they get sucked into the CG condensates and hence delivered to the SGs. As the SGs mature, presence of high zinc concentrations leads to hexamerization of insulin after its proteolysis. CGs can undergo aggregation and could represent one of the possible mechanisms for cargo segregation within the SGs.

The role of electrostatic interactions in mediating LLPS can be determined by testing the ability of proteins to form condensates at different salt concentrations. At high salt concentrations, protein-salt interactions dominate protein–protein interactions, thus inhibiting LLPS. Accordingly, when CGB was equilibrated at pH 6.1 in the presence of 500 mM NaCl (high salt), we did not observe condensate formation (Figure 8B), indicating that charge–charge interactions drive LLPS of CGB. To probe this further, all acidic residues in the five stretches were mutated to alanine (CGB_5(ED)/A) and their ability to undergo LLPS was monitored. CGB_5(ED)/A_GFP formed condensates in the absence of crowding agents, similar to the WT (Figure 8C). Since CGB does not have a well-defined calcium binding domain, we reasoned that these stretches are potential calcium-binding sites and tested whether CGB_5(ED)/A_GFP undergoes LLPS in the presence of calcium. Interestingly, condensate formation was not observed in the presence of 20 mM calcium, which otherwise induced robust LLPS of WT CGB (Figure 8D). These findings suggest that CGB LLPS is driven by electrostatic interactions and that the five stretches of acidic amino acids potentiate this reaction by binding to calcium.

## Discussion

To investigate the molecular role of CGs in proinsulin sorting, we purified fully functional recombinant CGA and CGB, which are significant drivers of granule biosynthesis, from mammalian cells. In combination with cell biology-based approaches, we describe a model for protein sorting towards SGs. We demonstrated that CGs undergo LLPS in conditions resembling the TGN milieu to form a client scaffold. The pH in the TGN lumen is the primary driver of CG condensation and client (proinsulin) recruitment, whereas calcium is dispensable for the reaction. Our experiments revealed that phase separated CGs act as a “client sponge,” incorporating clients based on their size. Finally, we demonstrated that LLPS is crucial for client recruitment. Solid aggregates of CGB fail to recruit soluble clients, and truncation of CGB, which significantly affects its capacity to undergo LLPS, impairs the formation of ectopic SG-like structures in non-secretory cells. These results are in contrast to the pre-existing dogma of cargo segregation between constitutive and regulated pathways (*21, 23, 38*)

### LLPS drives cargo condensation in the TGN

The predominant model that explains cargo sorting at the TGN in specialized secretory cells is termed “sorting for entry” or “sorting by aggregation” (*38*), and has two main tenets: (i) granin family proteins undergo aggregation at an acidic pH and in the presence of calcium, after which the resultant aggregate, surrounded by a membrane, buds off from the TGN to form the immature SG and (ii) granins aggregate and exclude proteins destined for lysosomes or constitutively secreted proteins, mediating cargo sorting towards the SG at the TGN. These conclusions were derived from experiments using isolated granule contents from adrenal or pituitary glands and purified CGA or CGB from vesicle lysates of bovine adrenal medullary chromaffin cells (*30, 35, 36*). Aggregation was monitored by either testing the abundance of natural soluble CGs in pellet fractions upon centrifugation or by changes in turbidity upon inducing aggregation at pH 5.5 and in the presence of high millimolar calcium concentrations. While informative, these studies utilized conditions that are not representative of the natural TGN environment, and thus have limited relevance in understanding how CG-dependent sorting at the TGN occurs in a physiological setting. Moreover, the molecular mechanism underlying the aggregation process as well as specific cargo sorting by the aggregate remain unclear. Our results show that it is the liquid-like state that is essential for cargo recruitment as the CGB aggregate fails to recruit client molecules. Thus, LLPS of CGs results in the formation of a scaffold within the TGN lumen that recruits soluble clients like proinsulin for their subsequent delivery to the SGs.

### In CG-LLPS, pH is the main driver while calcium is dispensable

In endocrine cells, the sorting process has been shown to strictly depend on the acidification of intracellular compartments because the addition of acidotropic drugs diverts regulated secretory proteins to the constitutive pathway, completely suppressing the formation of new secretory granules (*23*). CGs are synthesized in the ER and transported to the Golgi apparatus, where they are sorted and targeted to the SGs. Since LLPS of CGs facilitates cargo sorting towards the SGs, CGs must be in soluble form within the ER lumen and undergo LLPS at the TGN.

As proteins move along the secretory pathway, there is a pH gradient from an almost neutral pH (7.3) in the ER lumen that progressively becomes acidic from the *cis*- to the *trans*-Golgi, with a TGN pH of around 6.1 (*65–68*). Our results demonstrate that pH (6.1), resembling the TGN milieu is a critical driver of LLPS, whereas calcium is not necessary; this contrasts with previous studies that have indicated calcium is required for the aggregation of CGs. These results are further strengthened by our findings on the behavior of the CGB_5(ED)/A mutant protein, which underwent LLPS on its own at high protein concentrations but failed to do so at low protein concentrations in the presence of calcium. Notably, these results also highlight the five acidic stretches in CGB as potential calcium binding sites.

Similarly, LLPS of CGA in the presence of crowding agents was favored at pH 6.1, and no aggregation of CGA was observed in the presence of 40 mM calcium. These results further emphasize that acidic pH and not calcium is the most critical parameter governing the LLPS of CGs. It is not surprising considering that calcium concentrations in the ER (400 µM; (*40*)) are considerably higher than those at the TGN. We assume that protonation of some of negatively charged amino acid residues of CGA (PI 4.58) or CGB (PI 5.03) at low pH conditions can promote LLPS by reducing electrostatic repulsion; consequently, the acidic milieu could induce the condensation of CGs in the TGN.

### Differential effects of divalent cations on LLPS and aggregation of CGB

While calcium is not essential for LLPS of CGB, at concentrations above 5 mM, it boosts its ability to undergo LLPS. However, our results show that the effect is not limited to calcium as magnesium, manganese, and zinc can also induce LLPS. Manganese, which is highly abundant in the TGN, shows a higher potential than calcium for inducing LLPS based on the size and area covered by the droplets. The requirement of CGB to be present in a liquid-like scaffold within the TGN lumen is further emphasized based on the differences in cargo recruitment to condensates versus aggregates. LyzC, sorted to SG in INS1 832/13 cells and co-condensed with CBG in calcium-induced droplets, fails to be recruited to zinc-induced CGB aggregates. High zinc concentrations only occur in the SGs because of the localization of zinc transporters to the SGs (*22, 57*). Based on the differential effects of divalent cations on CGB and their location in the secretory pathway, we suggest that CGB undergoes a transition from a liquid-like condensate to an aggregate as it moves along the secretory pathway from the TGN to the SGs, and zinc-dependent aggregation may be required for client release within the SG. Interestingly, a recent preprint from the Hagen laboratory has demonstrated cargo segregation within secretory granules from the fly salivary gland in a calcium- and pH-dependent fashion that also relies on glycosylation (*69*). It remains possible that zinc plays an equivalent role in mammalian secretory cells.

### LLPS drives protein sequence-independent sorting at the TGN: A universal mechanism for receptor-independent sorting

According to the second tenet of the “sorting for entry” model, aggregation of CGs is thought to be a driving force at the TGN to mediate cargo sorting. Cargo molecules destined for SGs, such as proinsulin, neuropeptide Y (NPY), and processing enzymes PC1/3, PC2, and CPE, would be brought into the aggregate. In contrast, lysosomal hydrolases and other constitutive cargo molecules would be segregated from the aggregate, thus facilitating sorting. It remains unclear how molecules such as proinsulin are brought into the aggregate when there is no known receptor for this process and whether there is a specific sorting mechanism.

Our results challenge the specific sorting aspect of the “sorting by entry” model, as we show that ectopic expression of soluble proteins, such as LyzC-GFP or EqSOL-GFP, in INS1 832/13 cells are secreted constitutively in HeLa cells, resulting in their targeting to SGs. Interestingly, overexpression of CatD, a lysosomal hydrolase, also showed the same phenotype. Thus, we propose that phase-separated granin proteins form scaffolds that act like a sponge that brings soluble proteins together into the condensed phase. Secretion of LyzC-GFP in response to glucose stimulation in INS1 832/13 cells suggests that LyzC and insulin remain in SGs and are released together in response to glucose stimulation. These findings contradict the “sorting by retention” model (*70*) which proposes that non-secretory proteins are removed by vesicles that bud off from the SG, thus mediating differential sorting of regulated versus constitutively secreted cargo.

Given our data showing co-segregation of proinsulin, M6PR, and IGF2R within the TGN volume and routing of lysosomal hydrolases to SGs upon overexpression, we suggest that at endogenous levels, lysosomal hydrolases are trafficked to lysosomes in a cargo receptor-dependent manner. Upon overexpression, the receptors are likely to be saturated, and the hydrolases become clients of the CG scaffold. The expression of granin proteins drives the targeting of soluble proteins to SGs as the dominant pathway. In parallel, the significance of CGs in targeting clients to SGs is highlighted by the fact that expression of CGA reroutes LyzC to ectopic SGs. This is in line with previous findings showing that expression of *Xenopus laevis* CGA proteins in Cos7 cells directs co-expressed NPY to the ectopic CGA-derived granules (*71*). NPY, however, is targeted to SGs in specialized cells. Our results expand on this idea and show that CGA expression indeed reorganizes TGN sorting, as it can reroute LyzC to SGs, which is otherwise secreted constitutively from HeLa cells.

The nonspecific recruitment of clients to cells was in agreement with our *in vitro* assay findings. CGB scaffolds recruited LyzC, CatD, and a small positively charged polypeptide. However, it is interesting that large molecules such as IgG remain attached to the rims of CGB scaffolds and do not effectively enter the condensates, thus indicating a likely size-based cut-off for recruitment. Failure of IgG to co-precipitate along with granin aggregates in a previous study was used to define segregation of constitutive and regulated secretory cargoes, which formed the basis of sorting by aggregation (*72*). Our results refute the clear segregation of cargoes at the TGN based on either specific sequence or structural elements in the client proteins or their destination, and instead highlight that the larger size of IgG is what prevents it from being co-sorted to the SGs. SGs are not limited to pancreatic β-cells but are also present in the α cells of the pancreas, which secrete glucagon. We propose that instead of having unique receptors for different cargo molecules in these different cell types, cells utilize an LLPS sorting mechanism wherein cargo molecules are recruited to the granin sponge by their abundance within the TGN lumen, resulting in their targeting to SGs. Supporting this, the expression of GFP fused to a signal sequence directing it into the secretory pathway leads to its targeting to SGs in AtT-20 and PC-12 cells (*73*). Since molecules like insulin or NPY are crucial for physiology and metabolism, specialized cells like pancreatic β-cells express different members of the granin family, including CGA, CGB, SCG2, VGF. Our studies have demonstrated that both CGA and CGB can undergo LLPS, although with a difference in capacity, and *in silico* analysis of VGF also showed disordered regions within the proteins that may be indicative of LLPS. The capability of VGF to undergo LLPS needs to be tested, as ectopic expression of VGF also induces ectopic granule-like structures in NIH3T3 cells (*74*). Along similar lines, Spiess and colleagues showed that peptide hormone precursors including pro-vasopressin, pro-oxytocin, and POMC can form SGs in cell lines that normally lack regulated secretion (*62*). POMC, for instance, contains a highly disordered region, and it can be speculated that it forms similar scaffolds in the TGN that promote sorting of SG-targeted proteins. This suggests that the abundance of a phase separated scaffold protein in the TGN may regulate SG biosynthesis, which must be tested in future experiments.

Although client recruitment to condensates of synapsin or ZO1 seems to be specific, mediated by physical interactions between the scaffolds and the clients (*75, 76*), LLPS driven by IDRs of FUS, hnRNP A1, and elF4GII has been shown to recruit globular proteins to the condensates, presumably due to interaction with the IDR (*77*). We believe that such a mechanism, in combination with the size-based cut off, determines the composition of granin condensates.

While our studies explain how CGB scaffolds recruit clients for sorting to SGs, it remains unclear how the condensate is associated to the membrane for budding of the vesicles from the TGN. Former research has proposed that soluble SCGIII or CPE can act as a tether for CGA aggregates by binding to cholesterol (*78*). Based on our findings it is conceivable that granin scaffolds also accumulate tethering proteins such as SCGIII, thereby attaching the condensate to cholesterol-rich TGN membranes. Alternatively, phase separation of condensed protein complexes has been proposed to cause membrane budding and fission (*79, 80*). In this context, it was suggested that the assembly of IDPs artificially targeted to a membrane surface induces membrane curvature due to a substantial compressive effect in the membrane plane (*80–83*). It would indeed be interesting to test whether CG condensates can drive membrane budding using an artificial membrane and if tethering to the membrane occurs by cholesterol binding.

Overall, our findings identify LLPS as an essential cellular process that is responsible for cargo sorting of soluble proteins towards SGs and that failure of LLPS impairs SG biogenesis.

## Author Contributions

Conceptualization: AP, JVB

Methodology: AP, MT, YZ, SBS, JVB

Investigation: AP, MT, CKB, SCB, KER, JZ, BRR,

Visualization: AP, MT, CKB, SCB, KER, JZ, BRR,

Funding acquisition: JVB

Project administration: AP, JZ, SBS, JVB

Supervision: YZ, SBS, JVB

Writing – original draft: AP, JVB

Writing – review & editing: AP, MT, CKB, SCB, KER, JZ, BRR, YZ, SBS, JVB

## Acknowledgments

We highly appreciate constructive discussions with Christopher Burd, Xiaolei Su, Jonathan Bogan and James Rothman. We want to thank Cedric Asensio for the INS1 832/13 cells. We would like to thank Charlotte Ford and Mai Ly Tran for proofreading and discussion of the manuscript. Julia von Blume is funded by a Yale-start up grant and by the National Institute of General Medical Sciences of the United States National Institutes of Health under the award number GM134083-01, an Administrative Supplement 3R01GM134083-03S1 and a Project and Feasibility award from Yale Diabetic Research Center (GR112420). This work was also supported by startup funds provided by the Fraternal Order of Eagles Diabetes Research Center, University of Iowa to S.B.S., Department of Defense CDMRP grant PR190353 to S.B.S., and a National Institutes of Health Predoctoral Training Grant T32GM008629, PI Daniel Eberl to K.E.R.

