## Supplementary Figures and Materials and Methods for "Liquid-liquid phase separation facilitates the biogenesis of secretory storage granules"

### Supplementary Materials: Parchure et. al.

#### Materials and Methods

##### Cloning and Constructs

For cloning, all the PCR amplifications were done using Phusion High-Fidelity DNA polymerase, ligations using T4 DNA ligase which were both obtained from Thermo Fischer Scientific. Final vectors were also generated using Gibson assembly (New England Biolabs; (NEB)) and/or using Gateway assembly (Invitrogen) as and when indicated. All the restriction digestion was carried out utilizing restriction enzymes from NEB. All the reactions were carried out using manufacturer's protocols. Sequencing of the plasmids was carried out at Keck sequencing facility (Yale university).

For cloning of sfGFP and 6X-His tagged constructs of human CGA (pBT-PAF-CGA_sfGFP_6XHis) and CGB (pBT-PAF-CGB_sfGFP_6XHis), coding regions of CGA and CGB were cloned by PCR amplifications from expression plasmids from Origene (CGA; Cat# RC RC200492 and CGB; Cat# RC201744). Super-folder GFP (sfGFP) was cloned by PCR amplification from the RINS1 construct (*54*). The nucleotide sequence encoding for 6X-His was incorporated into the reverse primer for amplification of sfGFP and thus was after the sfGFP. pBT-PAF vector was linearized using NheI-HF and NotI-HF restriction enzymes (NEB). The two individual fragments encoding CGA/CGB and sfGFP were then inserted into the digested vector using Gibson assembly.

For generating truncation mutants of CGB (pBT-PAF-CGB_N term-sfGFP_6XHis, pBT-PAF-CGB_C term-sfGFP_6XHis and pBT-PAF-CGB_C term-mCherry_6XHis), the N terminus of CGB, encoding amino acids 1-334 of CGB, was cloned by PCR amplification from pBT-PAF-CGB_sfGFP_6XHis construct and the C terminus, compromising signal sequence and 335-667 of CGB, was ordered as a gBlock gene fragment from IDT. 6XHis-tagged sfGFP and mCherry were amplified from the RINS1 construct. Final vectors were generated using Gibson assembly to insert CGB_N/CGB_C and sfGFP/mCherry into the digested pBT-PAF vector.

For generating pBT-PAF-CGB_5(ED)/A_sfGFP_6XHis, a gBlock gene fragment was ordered from IDT where all the acidic residues (Glutamic acid and Aspartic acid) in the 5 stretches of CGB were mutated to Alanine. This gene fragment and 6XHis-tagged sfGFP amplified from the RINS1 construct were inserted into the digested pBT-PAF vector to generate the final construct using Gibson assembly.

pBT-PAF-proinsulin_mCherry_3XFLAG was generated in two steps. First, the full-length proinsulin gene was cloned from the RINS1 plasmid by excising out mCherry which is inserted within the C-peptide in this construct (*54*). For this purpose, the RINS1 plasmid was transformed dam$^-$/dcm$^-$ competent *E. coli* and digested by using ApaI enzyme to remove mCherry from C-peptide coding region. The bigger fragment was self-ligated and 16°C overnight using T4 ligase according to manufacturer's protocol. Derived plasmid was digested with NheI-HF and BamHI-HF enzymes and 374 bp fragment coding proinsulin was cloned into pBT-PAF vector linearized with NheI-HF and NotI-HF restriction enzymes (NEB) with 66 bp annealed fragment coding HA- and 6xHis-tags (pBT-PAF-proinsulin_HA_6XHis). To generate pBT-PAF-proinsulin_mcherry_3XFLAG, sequence encoding proinsulin was cloned from pBT-PAF-proinsulin_HA_6XHis and mCherry was cloned from RINS1 with the 3XFLAG sequence placed at the C-terminus of mCherry. The two fragments were then fused with overlap-extension PCR and digested with NheI-HF and NotI-HF and ligated with the digested pBT-PAF vector.

To generate pBT-PAF-6XHis_sfGFP, pBT-PAF-6XHis_sfGFP_Cab45 (*84*) was digested using AscI and NotI-HF restriction enzymes and was PCR amplified to introduce NheI and NotI sites. The fragment was double digested and then ligated into linearized pBT-PAF vector to obtain pBT-PAF_6XHis_sfGFP.

pLenti-LyzC_EGFP and pLenti-CatD_sfGFP were generated using gateway cloning. LyzC_EGFP was PCR amplified from pLPCX-LyzC_EGFP (*84*) by addition of appropriate nucleotides for gateway cloning reactions and cloned into the entry vector pDONR221 using the BP clonase reaction. LyzC_EGFP was cloned from the entry clone into the destination vector pLenti using LR clonase reaction to generate pLenti-LyzC_EGFP.

To generate pLenti-CatD_sfGFP, CatD fragment was PCR amplified from pBT-PAF-ssSUMO_CatD, by incorporating signal sequence from the CatD protein at the N-terminus and fused with sfGFP using extension overlap PCR also incorporating nucleotides for gateway reactions and the fragment was then cloned via sequential BP clonase and LR clonase reactions to obtain the destination vector.

Generation of recombinant adenovirus expressing CgB-CLIP has been previously described (*85*).

**Cell culture**

HEK293 and HeLa cells were maintained in DMEM high glucose (GIBCO) supplemented with 10% fetal bovine serum (FBS; GIBCO), 100 U/ml penicillin and 100 µg/ml streptomycin in 5% carbon dioxide at 37°C. Rat insulinoma INS 832/13 cells and INS 832/13 cells expressing SNAP tagged proinsulin or CatD-GFP and INS 832/13 cells dually expressing SNAP tagged proinsulin and LyzC-GFP were maintained in RPMI 1640 (GIBCO; Cat# 11879-020) containing 11 mM glucose and supplemented with 10% FBS, 100 U/ml penicillin and 100 µg/ml streptomycin, 10 mmol/l HEPES (americanBIO), 2 mmol/l Glutamax (GIBCO) , 1 mmol/l sodium pyruvate (GIBCO) and 50 µmol/l β-Mercaptoethanol (americanBIO) in 5% carbon dioxide at 37°C.

**Generation of stable cell lines for protein expression**

Doxycycline inducible HEK293 stably expressing GFP, and His tagged proteins or mCherry and FLAG tagged proinsulin proteins were generated using the transposon-based piggyBac system (*86*). HEK293 cells were transfected with a mixture of pBT-PAF, PB-RN and PBase (8:1:1 ratio) using Lipofectamine 2000 (Invitrogen) according to manufacturer’s protocol. Medium was exchanged after 4-6 hours. 48 hours post transfection cells stably expressing the plasmid cocktail were selected by treatment with the antibiotics puromycin dihydrochloride (10 µg/ml; Sigma-

Aldrich) and G418 disulfate salt (500 µg/ml; Sigma-Aldrich). Stable lines were then frozen down for subsequent use.

**Protein expression, purification of His-tagged proteins**

GFP/mCherry- and His-tagged proteins were purified from culture supernatants as described in Figure 2B using nickel-based column chromatography. Doxycycline-inducible HEK293 stably expressing GFP/mCherry- and His-tagged proteins were expanded in 10-20 15 cm dishes. Fully confluent cells were induced for protein expression by addition of doxycycline monohydrate (1 µg/ml; LKT Laboratories) in serum free DMEM high glucose also containing proteinase inhibitor aprotinin (1 µg/ml; Sigma-Aldrich) and antibiotic A23187 (1 µg/ml; Thermo Fisher Scientific Alfa Aesar) for 15-20 hours. Medium was collected and pre-cleared of cells by centrifugation and then by filtration using a 0.45 µm filter. The supernatant was then loaded onto a Ni-NTA column generated from cOmplete His-tag Purification Resin (Roche) which was equilibrated in Sodium Phosphate buffer, containing 500 mM NaCl; pH 6.8. The column was washed with the equilibrating buffer containing 10 mM imidazole and then eluted using the equilibrating buffer containing 250 mM imidazole. Imidazole was removed from the proteins by buffer exchanging the protein with Tri-HCl buffer containing 500 mM NaCl; pH 6.8 as well as 10% (vol/vol) glycerol and stored in it at -80°C after snap freezing the protein in batches in liquid nitrogen.

**Protein expression, purification of mCherry and FLAG tagged proinsulin**

mCherry- and FLAG- tagged proinsulin was purified from cell culture supernatants from HEK293 cells by expressing inducing the protein in cells grown in serum free DMEM high glucose medium containing doxycycline monohydrate (1 µg/ml) and aprotinin (1 µg/ml), which was loaded onto a column containing FLAG resin (Genscript; L00432-5). Protein was eluted using competitive elution with 3X FLAG peptide (APExBIO; A6001) and the eluted protein was buffer exchanged Tri-HCl buffer containing 500 mM NaCl; pH 6.8 also containing 10% (vol/vol) glycerol. Further purification was done using a round of size exclusion chromatography and the fractions containing protein of interest were flash frozen using liquid nitrogen and stored at -80°C till further use.

**Lentiviral production**

Lentiviruses were generated using HEK293 cells. HEK293 cells were plated on poly-Lysine coated 10 $cm^2$ dishes and grown in DMEM high glucose medium. The following day they were transfected with a plasmid cocktail containing packaging plasmid containing the gene of interest along with psPAX2 (Gag, Pol, Rev, and Tat), pMD2.G (VSV-G) using Lipofectamine 2000 according to manufacturer's protocol. On the next day, the medium was changed to INS 832/13 culture medium since the viruses were to be used for infecting INS 832/13. 48 hours post transfection medium containing virus particles was collected from cells and passed through a 0.45 μ filter and stored at 4°C. HEK293 cells were again supplemented with INS 832/13 culture medium for one more round of collection. 73 hours post transfection collection protocol was repeated and collection from two days was pooled and either used immediately for infection or stored at -80°C after aliquoting in batches.

**Generation of INS 832/13 stable cell lines**

Lentiviruses were used for infection of INS 832/13 cells for generation of stable cell lines expressing LyzC-EGFP and CatD-sfGFP. Stable cell lines were selected after antibiotic selection.

***In vitro* droplet formation assay**

Droplet formation of CGB-GFP without any crowding agents was monitored by microscopy after exchanging the storage buffer in which proteins were stored, containing high salt, to phase separation buffer, i.e., Tris-HCl; pH 6.1 containing 150 mM NaCl and 2.5% glycerol. 10 μl solution was plated on a coverslip glass bottom imaging dishes (Cellvis) and the droplets which were settled on the coverslip were imaged using Zeiss 880 confocal microscope by excitation using the 488 nm laser.

For inducing droplet/aggregate formation in presence of divalent cations, CGB solution was centrifuged at 4°C on a benchtop Sorvall Legend Micro 21R centrifuge at maximum speed for 20 mins to pre-clear of any existing droplets. Droplet formation was then induced by mixing 2.5 μM CGB-GFP (final concentration) with different concentrations of divalent cations in the phase separation buffer, followed by imaging of droplets which have been settled to the bottom of a coverslip glass bottom imaging dish.

For inducing CGB-GFP droplets in presence of crowding agents, 2.5 μM CGB-GFP (final concentration) was mixed with 1-3% PEG 8000 (Thermo Fischer scientific) and imaged on Zeiss 880 confocal microscope. Similarly, CGA-GFP droplets were induced using 3-5% PEG 8000 or in presence of 5% Dex500 (Sigma).

**Fluorescence recovery after photobleaching and analysis**

Partial FRAP experiments were carried out on settled droplets within 20-30 minutes of initiation of the experiment. FRAP module on the Zeiss 880 confocal microscope with a 63x oil objective (numerical aperture: 1.4) was used for the experiment. In all the experiments other than those described in Figure 3D, bleaching and subsequent imaging was performed using the excitation wavelength of 488 nm. A circular region of interest within the droplet was used for photobleaching with the laser operating at 100% capacity, and recovery of fluorescence within the bleached region was monitored using continuous imaging for next 1-1.5 minutes with the laser power less than 1% of its full capacity. For the experiment described in Figure 3D, a combination of 488 and 405 nm lasers were used for bleaching while recovery was monitored only by excitation at 488 nm. For analysis, the intensity profile in the bleached region was corrected using an unbleached region from other droplets in the image and the FRAP curves were then plotted using the corrected data sets.

**Effects of pH on droplet formation**

To test the effects of pH on droplet formation of CGB-GFP, CGB-GFP protein in storage buffer was exchanged to either phase separation buffer i.e., Tris-HCl; pH 6.1 containing 150 mM NaCl and 2.5% glycerol or in Tris-HCl; pH 7.3 containing 150 mM NaCl and 2.5% glycerol using Amicon concentrator (30kDa; 3x) and droplet formation was monitored by plating the protein solutions on a coverslip glass bottom imaging dish, followed by imaging on Zeiss 880 confocal microscope. To check for droplet formation at pH 5.2, CGB-GFP protein in storage buffer was exchanged to following buffer: Tris-HCl; pH 5.2 containing 150 mM NaCl.

To test the effects of pH on droplet formation of CGA-GFP, CGA-GFP protein in storage buffer was exchanged similarly to either phase separation buffer i.e., Tris-HCl; pH 6.1 containing 150 mM NaCl and 2.5% glycerol or in Tris-HCl; pH 7.3 containing 150 mM NaCl and 2.5% glycerol and droplet formation was induced in presence of 5% Dex500 and monitored by microscopic imaging.

**Effects of salt concentration of droplet formation of CGB-GFP**

To test the effects of pH on droplet formation of CGB-GFP, CGB-GFP protein in storage buffer was exchanged to either phase separation buffer i.e., Tris-HCl; pH 6.1 containing 150 mM NaCl and 2.5% glycerol or in Tris-HCl; pH 6.1 containing 500 mM NaCl and 2.5% glycerol (high salt) and protein solution was plated on a coverslip glass bottom imaging dish for imaging on Zeiss 880 confocal microscope.

**Client partitioning assay**

Client proteins were exchanged from their storage buffers into phase separation buffer. Peptides were dissolved into phase separation buffer to a final concentration of 10 μM. CGB-GFP and clients were diluted to a final concentration of 2.5 μM and 1 μM and droplet/aggregate formation was induced using either PEG 8000 or 20 mM calcium (droplets) or 20 mM zinc (aggregates). Droplets were imaged using Zeiss 880 microscope.

For quantifying recruitment of Cy3-LyzC to condensates or aggregates, imaging was performed under same settings in both GFP and the Cy3 channel. Region of interest were drawn using the GFP channel (CGB-GFP) in imageJ to measure the intensity in the GFP channel and in the Cy3 channel. Data was represented as ratio of signal in Cy3 channel to that in the GFP channel and compared in calcium induced and zinc induced conditions. Similar methodology was used to measure the recruitment of Cy3-CatD and Cy3-polyD to CGB-GFP condensates.

**Protein detection by Immunoblotting, Coomassie staining and Immunofluorescence**

For western blotting, proteins were transferred from SDS gels to a nitrocellulose membrane using web blot system from Bio-Rad laboratories. After transfer to membranes, they were incubated with 5% milk made in Tris buffered saline containing 0.1% Tween20 for at least one hour. Membranes were incubated with specific primary (overnight incubation) and HRP-coupled secondary antibodies (one hour incubation) and proteins were detected using chemiluminescence (Thermo Fischer Scientific) using ChemiDoc imaging system (Bio-Rad laboratories).

For detecting protein using Coomassie staining, SDS gels were rinsed with water following which stained using 0.1% Coomassie Brilliant Blue solution made in 40% vol/vol methanol and 10% vol/vol acetic acid and destained using 40 % vol/vol methanol and 10% vol/vol acetic acid solution and stored in water prior to imaging.

For immunofluorescence-based detection of proteins within the TGN volume, 832/3 cells were plated on HTB9 coated coverslips at low density and cultured overnight as previously described (*29*). Cells were fixed in 10 % neutral-buffered formalin and incubated overnight with primary antibodies as indicated. Highly cross-adsorbed fluorescent conjugated secondary antibodies (whole IgG, donkey anti-guinea pig-AlexaFluor 488, donkey anti-rabbit rhodamine red-X, donkey anti-mouse AlexaFluor 647; Jackson ImmunoResearch) were used for detection. Cells were counterstained with DAPI and mounted using Fluorosave (Calbiochem).

For detection of proteins post Brefeldin A treatment, Cells were then fixed using 4% paraformaldehyde made in PHEM buffer and then permeabilized using PHEM buffer containing 0.3% NP-40 and 0.05% Triton X-100 for 5 minutes. Primary and secondary antibodies were diluted in PHEM buffer containing 0.05% NP-40, 0.05% Triton X-100 and 5% serum. Coverslips were mounted using Prolong Gold also containing DAPI (Thermo Fischer Scientific) and imaged using Zeiss 880 confocal microscope.

For all other experiments, cells were either grown on coverslips or glass bottom dishes (Cellvis) and fixed using 4% paraformaldehyde (Electron Microscopy Sciences) made in phosphate buffered saline (PBS) and permeabilized using 0.4% saponin (Sigma) made in PBS containing 4% BSA (americanBIO) for at least one hour. Cells were stained in respective primary antibodies and fluorophore conjugated secondary antibodies (Invitrogen). Coverslips were mounted using Prolong Gold also containing DAPI and imaged using Zeiss 880 confocal microscope.

**Pulse chase experiments**

For SNAP-tag and CLIP-tag labeling, 832/3 cells stably expressing proCpepSNAP (*29*) were treated with recombinant adenovirus expressing CgB-CLIP and pulse-chase labeled as previously described (*85*) for the indicated times.

**3D rendering of TGN volume**

Images of INS 832/13 stained with antibodies to TGN38 and soluble cargo or receptors were captured on a Leica SP8 confocal microscope using a HC PL APO CS2 40x/1.40 oil objective with 5x zoom as z-stacks (5 per set, 0.3 µm step, 0.88 µm optical section) and deconvolved (Huygen's Professional). Golgi volume mask was created using surface rendering of the Golgi identified by TGN38 immunostaining (Imaris, Bitplane) and line intensity profile of immunostaining within the Golgi was generated using ImageJ software.

**Electron microscopy**

C57BLKS/J mice were purchased from Jackson Laboratories and maintained as an active breeding colony. Islets were isolated from 14-week-old male mice via collagenase V digestion and purified using Histopaque 1077 and 1119. Islets were allowed to recover overnight in RPMI supplemented with 10% fetal bovine serum and 1 % penicillin and streptomycin and maintained at 37°C in 5% $CO_2$. All animal protocols were approved by the University of Iowa Institutional Animal Use and Care Committee. Isolated islets were washed by PBS and fixed in 2.5% glutaraldehyde and 4% paraformaldehyde at 4°C overnight. Islets were washed with 0.1 M sodium cacodylate three times and post-fixed in freshly made 1% (w/v) reduced OsO4 and 1.5%

(w/v) cyanoferrate in 0.1 M sodium cacodylate for 1 h. Islets were rinsed three times with water and processed for successive dehydration (50% ethanol, 2 × 5 min; 70% ethanol, 2 × 5 min, 90% ethanol, 2 × 5 min; 100% ethanol, 3 × 15 min; and propylene oxide, 15 min). Infiltration was performed using Epon-Mix, and samples were incubated at 60°C overnight for polymerization as previously described (*87*, *88*). Resin blocks were cut to ultrathin (50-70 nm) sections with a diamond knife and mounted on Formvar-coated copper grids. Grids were double contrasted with 2% uranyl acetate and then with lead citrate at room temperature and washed immediately with excessive water. Images were captured at 8,000x and 12,000x magnifications by a JEOL JEM-1400 transmission electron microscope. All EM-related reagents were from Electron Microscopy Sciences (EMS; Hatfield, PA).

**Imaging of ectopic granules in HEK 293 cells**

HEK293 cells stably expressing CGB-GFP and truncation mutants of CGB were plated on glass bottom imaging dishes from Cellvis. Cells were induced from protein expression by addition of doxycycline monohydrate (1 µg/ml; LKT Laboratories) for 10 hours following which cells were fixed using 4% paraformaldehyde (Electron Microscopy Sciences) and co-stained with DAPI to label the nuclei. Cells were imaged on the Zeiss 880 confocal microscope. The bottom plane closest to the coverslip was imaged using a 63x oil objective (numerical aperture: 1.4). In some cells expressing GFP tagged C-terminal portion of CGB, we observed signal from the nucleus and these cells were excluded from imaging.

**Labeling of CatD**

Recombinant human CatD protein was purchased from Abcam. It was labeled with Cy3 using a kit from AAT Bioquest according to manufacturer's protocols.

**Secretion Assays**

Secretion of LyzC-EGFP in response to glucose stimulation was monitored from cells stably expressing LyzC-EGFP and SNAP tagged proinsulin. Cells were plated in a 10 $cm^2$ dish and assayed at greater than 80% confluency. On the previous day of the assay, cells were incubated in INS 832/13 containing medium but with 5 mM glucose concentration instead of the regular 11 mM glucose concentration. After overnight incubation, cells were incubated in secretion assay buffer as described in (*61*) for 2 hours followed by incubation in either serum free INS 832/13 medium with either 3 mM glucose (basal) or 15 mM glucose and 35 mM KCl (stimulated) for 2 hours. Medium was collected after centrifugation at 800 × g for 5 minutes for removal of any floating cells and then concentrated using Amicon concentrators (3kDa cut off). Levels of proteins in supernatants and cell lysates were analyzed using western blotting and probed using α-GFP antibody to detect the levels of LyzC-GFP and α-SNAP antibody to monitor the secretion of SNAP tagged C-peptide, a proxy for insulin secretion. Intensity of bands in cell lysates and supernatants were measured using densitometry and amount of LyzC-GFP protein in the supernatant was normalized to the levels in cell lysates in both basal and stimulated conditions.

**Golgi ministacks**

INS 832/13 cells were plated on glass coverslips and treated with 33 μM Nocodazole (Santa Cruz) for three hours, following which cells were fixed using 4% paraformaldehyde and processed for immunofluorescence. Control cells were treated with 0.1% DMSO.

**BFA treatment**

INS 832/13 cells were plated on glass coverslips and treated with Brefeldin A (5μg/ml; Cell Signaling Technology) for three hours at 37°C. Control cells were treated with 0.05% DMSO.

**Antisera**

Antibodies used for immunofluorescence were as follows: Insulin (Thermo Fischer Scientific; MA1-10517 and Dako; A0564), CGB (Synaptic systems; 259 10 and Proteintech, 14968-1-AP), CGA (Novus biologicals; NB120-15160), GM130 (BD Transduction; 610822), TGN 38 (Novus Biologicals NB300-575 and BD Biosciences; 610898), CPE (Proteintech; 13710-1-AP) M6PR (Proteintech; 16795-1-AP), IGF2R (Proteintech; 20253-1-AP), SCGIII (Proteintech; 10954-1-AP), PCSK2 (Invitrogen; PA1-058), HA (Roche/Sigma Aldrich; 11867423001) Cathepsin B (R&D

systems; AF965). Antibody against Cab45 is a custom generated antibody as described previously (*89*).

For purposes of western blotting antibodies against GFP (Roche/Sigma Aldrich; 11814460001) SNAP-tag (NEB; P9310S) and β-actin (Sigma Aldrich; A5441) were used.

**Quantification and statistical analysis**

Fiji was used for analyzing images from *in vitro* phase separation assays. For each image, droplet particles with size larger than 0.01 $um^2$ were thresholded and selected. The selection was applied to the original images for measurement. For each image, the size of droplets was measured, and sum of droplet sizes was divided by the area of a field of view to obtain the area coverage of condensates. The distribution frequency of droplets or condensate coverage from a set of images was plotted using Prism 9.

Fiji and in-built measurement were used for analyzing images of ectopic granules in HEK cells. To quantify the area covered by granules in cell, each cell body outlined by peripheral GFP signals was captured as an ROI. For each ROI, GFP signals from cytosolic granules were measured and signals from the nucleus (stained with DAPI) or Golgi were excluded. Area covered by granules was divided by total area of an ROI and area coverage from a set of images was plotted using Prism 9.

#### Supplementary Figure legends

**Supplementary Figure 1**

**A)** Images obtained from INS 832/13 cells which were stained with TGN38 (red) and PC (green), treated with either 0.05% DMSO (control; top) or 5 µg/ml Brefeldin A (bottom). Shown here is a single slice from a confocal stack. Note the disassembly of TGN38 based on staining upon Brefeldin A treatment which is accompanied by a loss of PC2 puncta in the perinuclear region, marked using dashed line. **B)** Single slice from a confocal stack obtained by imaging INS 832/13 cells stained using GM130 (red) or PC2 (green) upon treatment with 0.1% DMSO or 33 µM nocadazole. Note the localization of PC2 puncta in vicinity of GM130 signal, which is dispersed throughout the cell upon nocadozole treatment, marked using arrows. This is in stark contrast to the perinuclear distribution of PC2 puncta in control cells.

**Supplementary Figure 2**

**A)** Protein sequence of the super-folder GFP with the momomerizing mutation which was used for generation of fusion constructs used for studying LLPS. R (arginine) at position 206 is highlighted in red, which is present in place of valine or alanine in parental super-folder GFP protein. **B)** A panel of images monitoring recovery of fluorescence of CGA-GFP droplets, induced by addition of 5% Dextran, after bleaching a small region within the droplets. **C)** Graph quantifying the fluorescence recovery in time. Data represented as mean +/- s.d. (error blanket) from six independent droplets.

**Supplementary Figure 3**

**A)** Images obtained from plating a solution of CGB-GFP (2.5 µM final concentration) to monitor the presence or absence on liquid like condensates either without or with 250 µM or 5 mM calcium. Note that droplet formation is induced only in presence of 5 mM calcium. **B)** Images obtained from plating a solution of CGB-GFP (2.5 µM final concentration) in presence of 5 mM magnesium, calcium, or manganese respectively. While manganese and calcium can induce droplets at this concentration, magnesium fails to do so. **C)** Images obtained from plating a solution of CGA-GFP (2.5 μM final concentration) in presence of either 250 μM, 20 mM or 40 mM calcium to test for the presence or absence of droplet formation. No droplets are seen event at 40 mM which is the highest calcium concentration.

**Supplementary Figure 4**

**A)** Confocal images obtained from INS 832/13 cells stained with proinsulin (green), CPE (red; left), M6PR (red; middle), IGF2R (red; right) and TGN38 (magenta; surface rendered). TGN38 staining was used for obtaining the Golgi mask. These are the cells which have been shown Figure 5A-C. **B)** Single slice from a confocal image obtained from INS 832/13 cells expressing mCherry and Apex tagged proinsulin (red) and stained with an antibody against Cathepsin B (green). Area in white denotes the Golgi region which was marked using GM130 staining which is not shown here.

**Supplementary Figure 5**

**A)** Images obtained by plating 10 μM of either CGB-GFP or CGB_Nterm-GFP or CGB_Cterm-mCherry without any crowding agent. Upon equilibration at pH 6.1 Note that droplet formation is seen only in CGB-GFP solution under these conditions. **B)** Images obtained by plating of 2 μM of CGB-GFP or CGB_Nterm-GFP or CGB_Cterm-GFP in presence of 3% PEG8000 to monitor droplet formation in these conditions. Only few small droplets are observed in CGB_Cterm-GFP as compared to CGB-GFP or CGB_Nterm-GFP. **C)** Western blots to compare the expression levels of CGB_Nterm-GFP and CGB_Cterm-GFP from HEK293 cells stably expressing these constructs and probed using GFP antibody (top). B-actin (bottom) was used as a loading control.

**Supp Fig1**

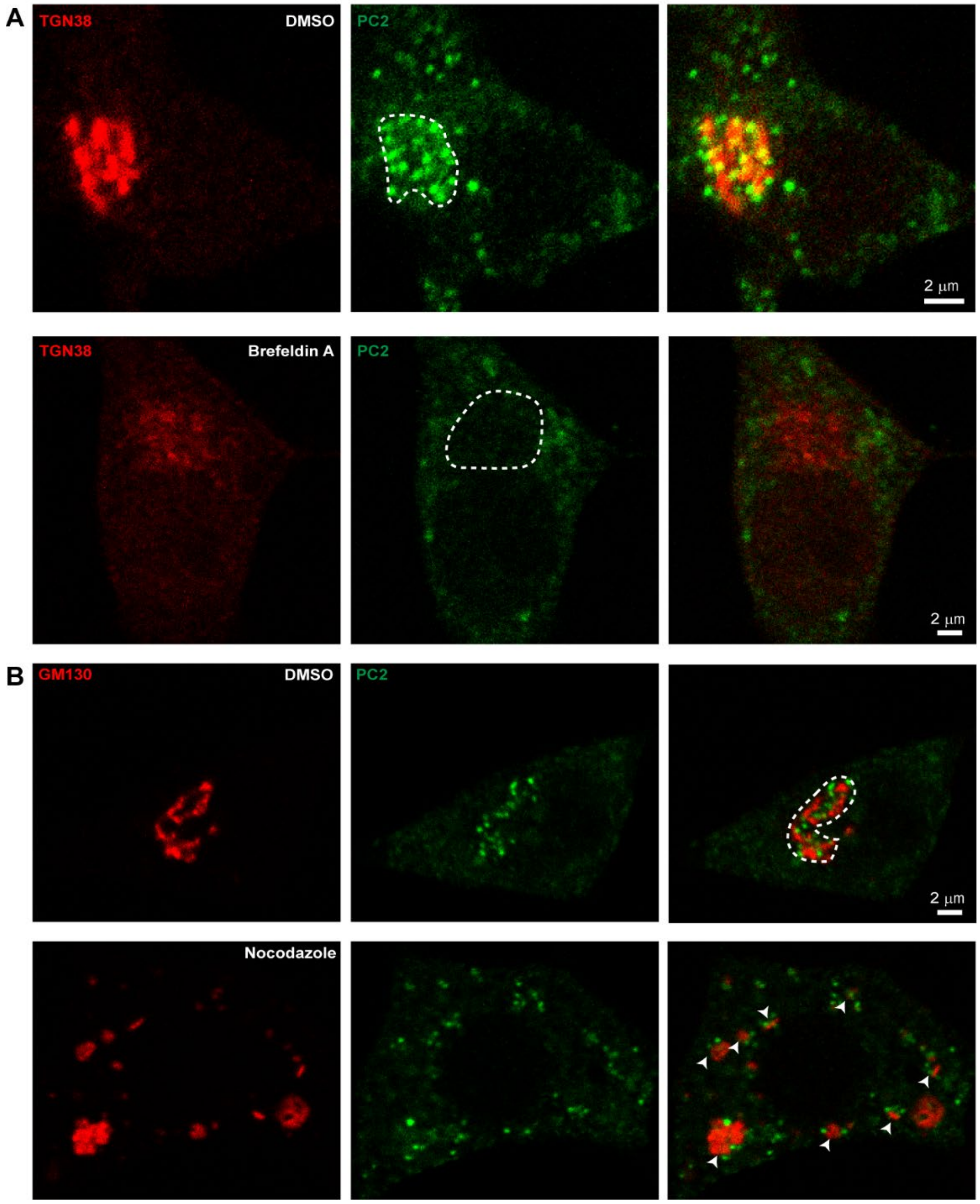
A
TGN38
DMSO
PC2
2 μm
TGN38
Brefeldin A
PC2
2 μm
B
GM130
DMSO
PC2
2 μm
Nocodazole

**Supp Fig2**

**A** sf-GFP protein sequence

| | | |
|---|---|---|
| **1** | MSKGEELFTG VVPILVELDG DVNGHKFSVR GEGEGDATNG KLTLKFICTT GKLPVPWPTL | **60** |
| **61** | VTTLTYGVQC FSRYPDHMKR HDFFKSAMPE GYVQERTISF KDDGTYKTRA EVKFEGDTLV | **120** |
| **121** | NRIELKGIDF KEDGNILGHK LEYNFNSHNV YITADKQKNG IKANFKIRHN VEDGSVQLAD | **180** |
| **181** | HYQQNTPIGD GPVLLPDNHY LSTQSRLSKD PNEKRDHMVL LEFVTAAGIT HGMDELYK | **238** |

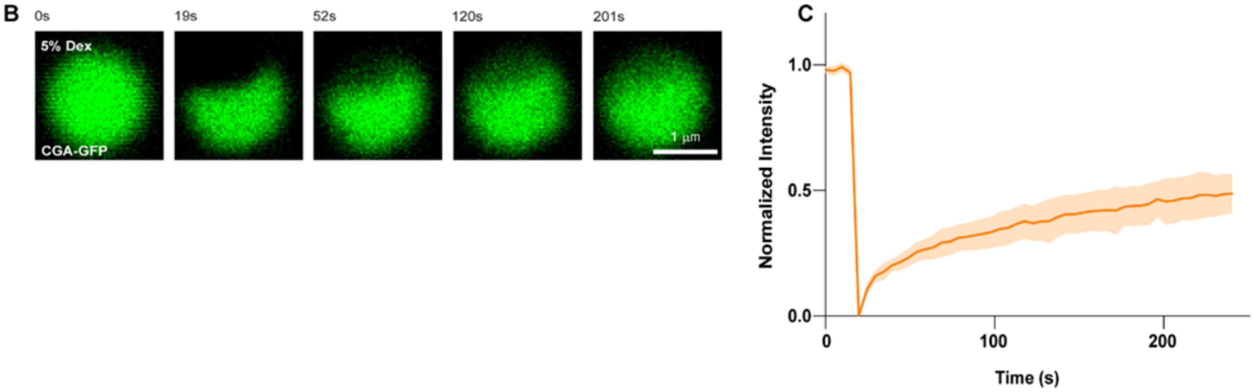

#### Supp Fig3

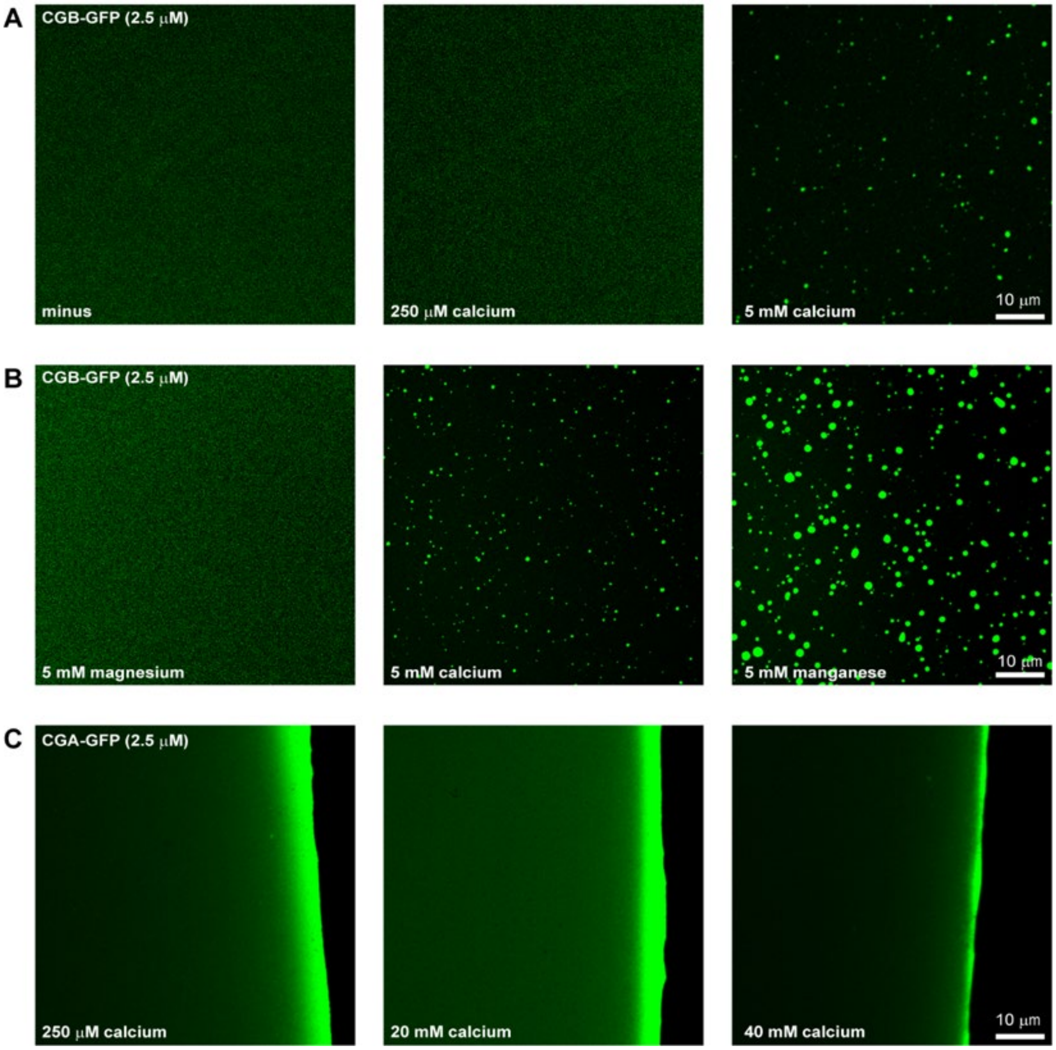

A
CGB-GFP (2.5 μM)
minus
250 μM calcium
5 mM calcium
10 μm
B
CGB-GFP (2.5 μM)
5 mM magnesium
5 mM calcium
5 mM manganese
10 μm
C
CGA-GFP (2.5 μM)
250 μM calcium
20 mM calcium
40 mM calcium
10 μm

#### Supp Fig4

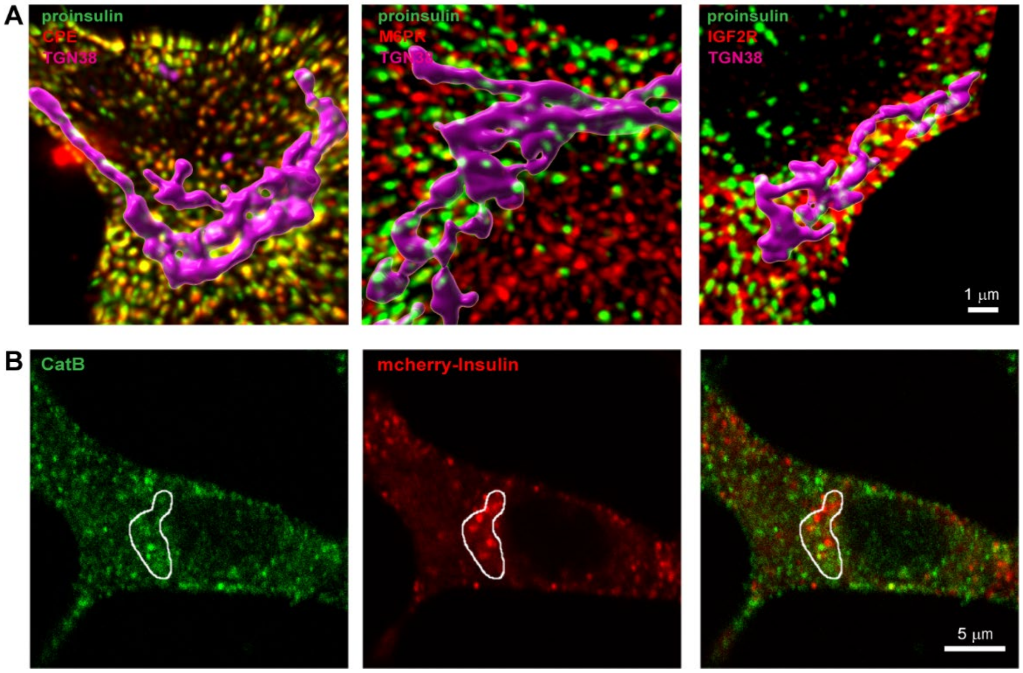
A
proinsulin
CPE
TGN38
proinsulin
M6PR
proinsulin
IGF2R
TGN38
1 μm
B
CatB
mcherry-Insulin
5 μm

**Supp Fig5**

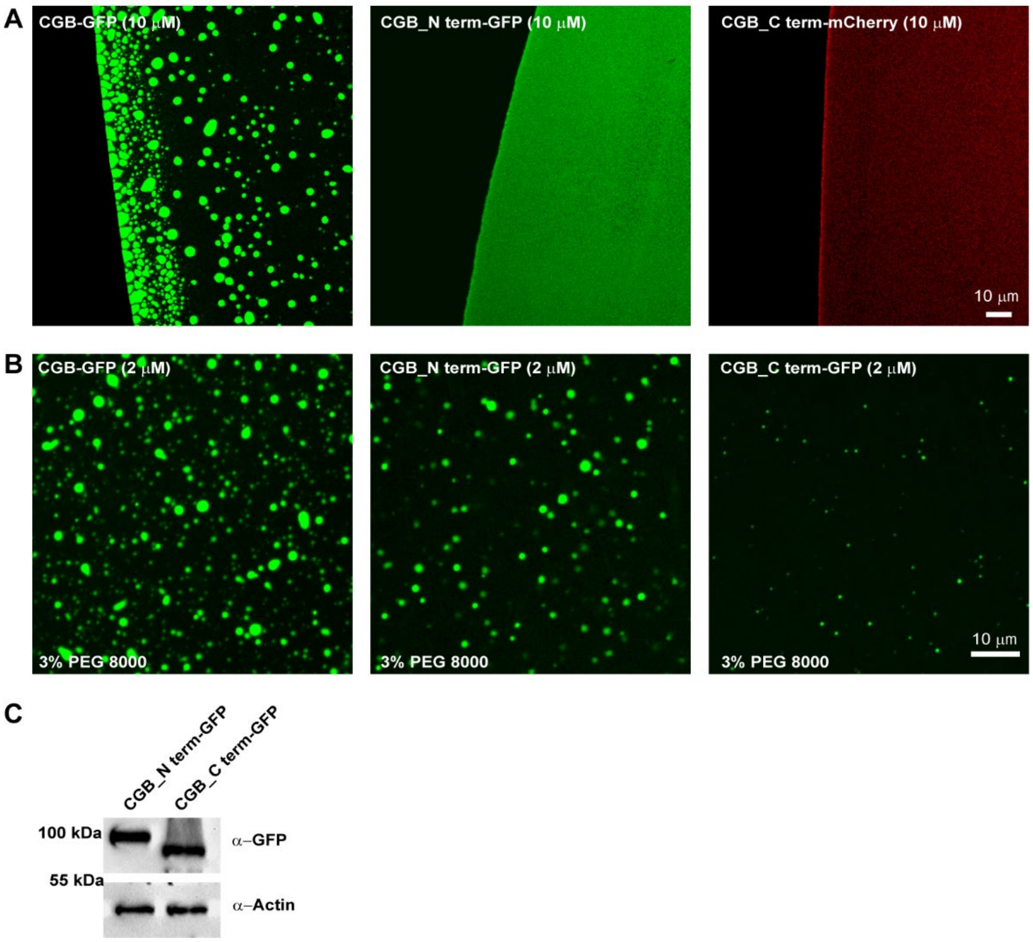

A
CGB-GFP (10 μM)
CGB_N term-GFP (10 μM)
CGB_C term-mCherry (10 μM)
10 μm
B
CGB-GFP (2 μM)
CGB_N term-GFP (2 μM)
CGB_C term-GFP (2 μM)
3% PEG 8000
3% PEG 8000
3% PEG 8000
10 μm
C
CGB_N term-GFP
CGB_C term-GFP
100 kDa
55 kDa
α–GFP
α–Actin
